# LCK-14-3-3ζ-TRPM8 axis for regulating TRPM8 function/assembly promotes pancreatic cancer malignancy

**DOI:** 10.1101/2022.01.26.477835

**Authors:** Yuan Huang, Shi Li, Zhijie Wang, Qinfeng Liu, Shunyao Li, Lei Liu, Weiwei Zhao, Kai Wang, Rui Zhang, Declan Ali, Marek Michalak, Xing-Zhen Chen, Cefan Zhou, Jingfeng Tang

## Abstract

The transient receptor potential melastatin 8 (TRPM8), function as a Ca^2+^-permeable channel in the plasma membrane (PM). Dysfunction of TRPM8 is associated with human pancreatic cancer and several other diseases in clinical patients, but with unclear underlying mechanisms. Here, we found lymphocyte-specific protein tyrosine kinase (LCK) directly interacts with TRPM8 and potentiates TRPM8 phosphorylation at Y1022. LCK positively regulated channel function characterized by increased TRPM8 currents densities through enhancing TRPM8 multimerization. Furthermore, 14-3-3ζ interacted with TRPM8 and positively modulated channel multimerization. LCK significantly enhanced the binding of 14-3-3ζ and TRPM8, whereas mutant TRPM8-Y1022F impaired TRPM8 multimerization and the binding of TRPM8 and 14-3-3ζ. Knockdown of 14-3-3ζ impaired the regulation of LCK on TRPM8 multimerization. Additionally, TRPM8 phosphotyrosine at Y1022 feedback regulated LCK activity by inhibition of Tyr505 phosphorylation and modulation of LCK ubiquitination. Finally, we revealed the importance of TRPM8 phosphorylation at Y1022 in the proliferation, migration and tumorigenesis of pancreatic cancer cells. Our findings demonstrate that LCK-14-3-3ζ-TRPM8 axis for regulating TRPM8 assembly, channel function, LCK activity and providing potential therapeutic targets for pancreatic cancer.

## Introduction

Pancreatic cancer is an aggressive malignancy with high mortality and only 5-7% of patients living longer than five years after diagnosis (Vincent et al, 2011). Although recent advances in radiotherapy and chemotherapy have been shown promising results, the overall prognosis and survival rates of pancreatic cancer patients are limited (Kobi et al, 2020). Therefore, understanding the biology of pancreatic cancer and identifying putative therapeutic targets in clinical treatment are urgently in need. Transient receptor potential melastatin 8 (TRPM8), the first identification of a prostate-specific gene, was functionally characterized as a cold receptor owing to its activation by cold temperature and substances that mimic cold sensation such as menthol and icilin, and plays a central role in thermosensation (McKemy et al, 2002). Recently, several studies revealed that TRPM8 exhibits aberrant expression and contributes to the development and progression of pancreatic cancer (Yee, 2016; Yee et al, 2012). Identification of the mechanisms by which TRPM8 mediate its biological functions is expected to develop into a molecular biomarker and therapeutic target in pancreatic cancer.

TRPM8 belonging to the TRP channels family, functions as a nonselective, voltage-gated, and Ca^2+^-permeable channel, which must be correctly expressed and assembled in the plasma membrane (PM). Previous studies showed that four monomers were assembled to form a homologous tetramer of functional TRPM8 channels (Erler et al, 2006; Phelps & Gaudet, 2007; Tsuruda et al, 2006). Although the C-terminal coiled coil (K1066-K1104) of TRPM8 has been implicated in channels multimerization (Erler et al, 2006; Phelps & Gaudet, 2007; Tsuruda et al, 2006), the mechanism remains obscure. Additionally, several molecules such as PIP2, PKA, PKC, TRP channel associated factors (TACF1 and TACF2), tripartite motif-containing 4 (TRIM4) and ubiquitin-like modifier activating enzyme 1 (UBA1) have been demonstrated to modulate TRPM8 channel expression in PM and activity by us and others (Abe et al, 2006; Asuthkar et al, 2015; Gkika et al, 2015; Huang et al, 2021; Rohacs et al, 2005; Zheng et al, 2018). Earlier studies revealed that 4-amino-5-(4-chlorophenyl)-7-(dimethylethyl) pyrazolo[3,4-d] pyrimidine (PP2, a selective Src family tyrosine kinase inhibitor) inhibited TRPM8 function in SH-SY5Y and HEK293T cells and TRPM8 is phosphotyrosined by Src, a membrane of nonreceptor Src family kinases, and partly by a representative of receptor PTKs, TrkA without identification of the exact site(s) (Manolache et al, 2020; Morgan et al, 2014). Thus, the mechanism of TRPM8 regulated by tyrosine kinases needs to be further investigated.

The lymphocyte-specific protein tyrosine kinase (LCK), which function as a Src-related nonreceptor protein tyrosine kinase, has emerged as one of the key molecules regulating T-cell functions (Adler et al, 1988; Voronova & Sefton, 1986). The dysregulated LCK, like that of other Src kinases, is associated with various diseased conditions like cancers, asthma, diabetes, etc (Kumar et al, 2018). The mechanistic insights into the regulation of LCK activity is sophisticated. Currently, LCK activity is predominantly regulated via reversible and dynamic phosphorylation of two tyrosine residues, one within the “activation loop” of the catalytic domain, Y394, and the other at the carboxy-terminus (C-terminus) of the protein, Y505 (Eck et al, 1994; Peri et al, 1993; Yamaguchi & Hendrickson, 1996). Blocking phosphorylation on Tyr394 (Y394F) largely reduced LCK activity, whereas inhibition of Tyr505 phosphorylation (Y505F) stimulated LCK activity (Eck et al, 1994; Peri et al, 1993; Yamaguchi & Hendrickson, 1996). In addition to phosphorylation, ubiquitination is also involved in regulating LCK activity (Choi et al, 2010; Giannini & Bijlmakers, 2004; Rao et al, 2002). For example, heat shock protein 90 (Hsp90) prevents the active form Y505F of mutant LCK from being targeted for degradation by ubiquitination (Giannini & Bijlmakers, 2004). 14-3-3 is a family of small acidic proteins that widely expressed in many organisms and tissues, and consists of seven highly conserved ~30 kDa isoforms (β, ε, γ, η, σ, τ, ζ) (Fu et al, 2000). By forming dimers, 14-3-3 predominantly bind phosphorylated proteins to modulate their targets at various levels, such as subcellular localization, stability, multimerization, phosphorylation, biological activity, or dynamic interactions (Gardino et al, 2006; Liu et al, 2021). TRPM7, belong to TRP channels family, binds to 14-3-3 for modulating channel cellular localization that requires autophosphorylation at S1403 (Cai et al, 2018). Apart from TRPM7, 14-3-3 involving in the regulation of other TRP channels function is limited. In addition, there is no report that 14-3-3 is involved in the regulation of LCK on target proteins.

In this study, we employed a variety of biological approaches to identify and unveil the mechanisms of LCK-14-3-3ζ-TRPM8 axis on regulating TRPM8 assembly, channel function and LCK activity and provided the importance of TRPM8 phosphotyrosine at Y1022 on the pancreatic cancer cells, which may be potential therapeutic targets for pancreatic cancer.

## Results

### LCK-TRPM8 interaction for positively modulating TRPM8 phosphotyrosine

We have previously reported GST pull-down assay in combination with mass spectrometric (MS) analysis to screen candidate proteins from a ~60 kD band that bind to the C-terminus of TRPM8 (M8C, amino acid 980-1104). The Ub-ligase E3 for TRIM4 was identified as a novel partner of TRPM8 by our earlier report (Huang et al, 2021). Within the same screen, we also identified peptides for BLK, LCK, LYN, belonging to Src family kinases, which were found in a similar size of ~60 kD band using co-immunoprecipitation (Co-IP) and MS assays in MCF7 cells (**Fig. *EV*1, *A-D***).

To further document the interaction between TRPM8 and these Src family kinases, we performed an *in vitro* GST pull-down assay. The results indicated that purified GST-M8C more efficiently pulled down HA tagged LCK compared with HA tagged BLK, and LYN from HEK293T cell lysate (**Fig. 1*A***). Moreover, we also confirmed that the BLK, LCK, and LYN kinases interacted with full-length TRPM8 and the LCK-TRPM8 binding was obviously strongest (**Fig. 1*B***). In consist with the above assays, reciprocal Co-IP experiments showed that BLK, LCK, LYN respectively associated with C-termini of TRPM8 in a protein complex, and the strongest interaction of C-termini of TRPM8 and LCK was clearly observed (**Fig. 1*C***.

**Figure 1.**
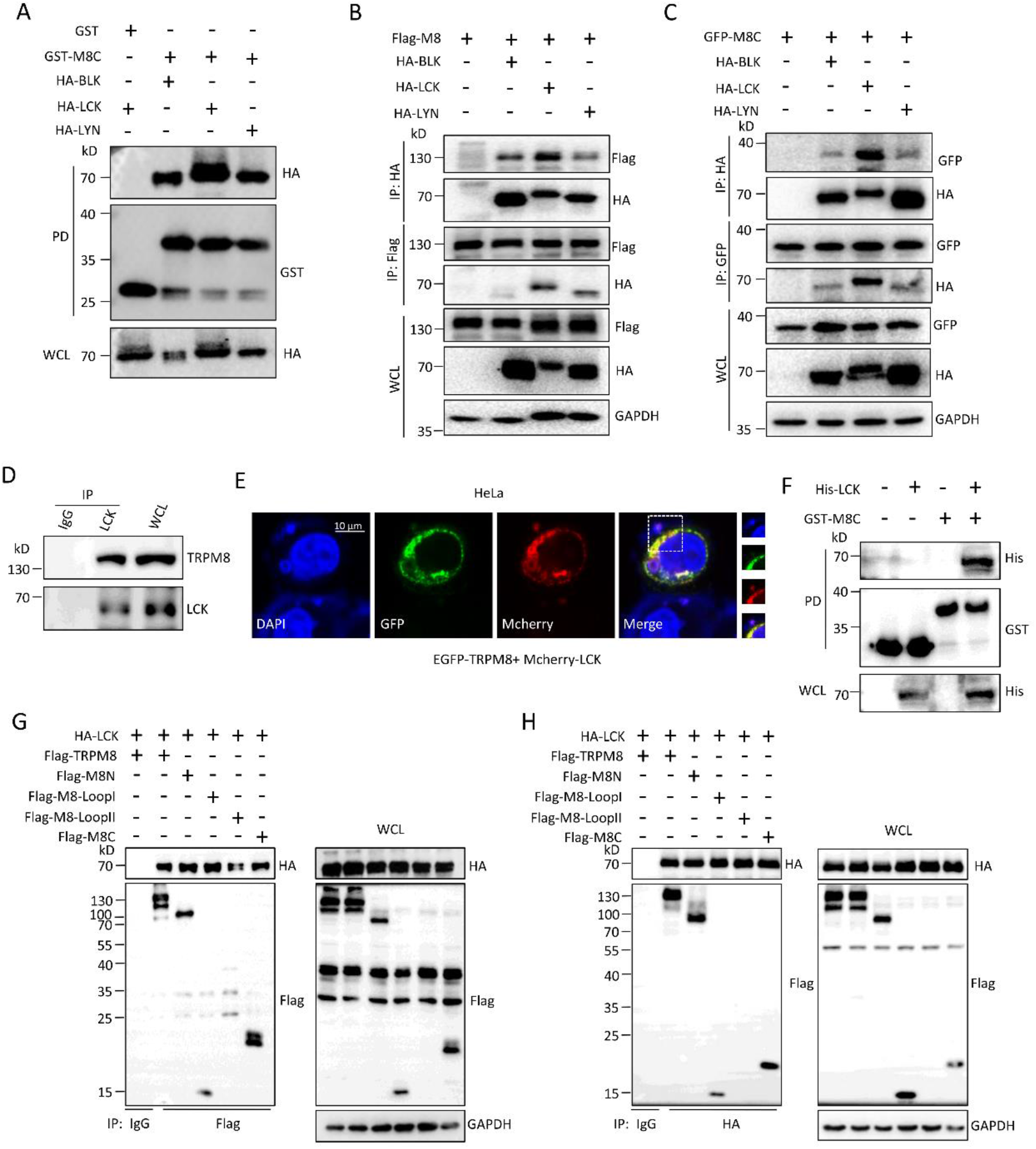
Src family kinases for BLK, LCK, and LYN interaction with TRPM8. (**A**) GST pull-down assays to assess the interaction between purified GST-tagged C-terminus of TRPM8 (GST-M8C) expressing in *E.coli* BL21 bacteria and different HA-tagged Src family kinases expressing in HEK293T cells. Western blotting (WB) was performed using the indicated antibodies. (**B-D**) Co-immunoprecipitation (Co-IP) assays. (**B**) HeLa cells were transfected with the indicated constructs along with Flag-tagged full-length TRPM8 (Flag-TRPM8). Immunoprecipitation (IP) was performed with an anti-HA or anti-Flag antibody, and the samples were analyzed by immunoblotting with the indicated antibodies. (**C**) Similar Co-IP in ***B*** but with protein extracts from HeLa cells co-transfected with the indicated constructs along with GFP-tagged C-terminus of TRPM8 (GFP-M8C). (**D**) Co-IP as in ***B*** and ***C*** but with protein extracts from native PANC-1 cells. (**E**) Representative confocal imaging of co-localization of mcherry-LCK and GFP-TRPM8 in HeLa cells. Overlay images show co-localization of green signals (TRPM8) and red signals (LCK), which generated yellow signals in HeLa. Nuclei were stained with DAPI (blue). Scale bars, 10 μm. (**F**) Assay of the interaction *in vitro* between purified His-LCK fusion and GST-tagged C-terminus of TRPM8 (GST-M8C) from *E.coli* bacteria. (**G-H**) HEK293T cells co-expressing HA-LCK constructs with a series of mutant Flag-tagged cytoplasmic domain of TRPM8 were harvested for Co-IP assays. All studies were repeated at least three times. GFP, green fluorescent protein. All studies were repeated at least three times.

Src family kinases are involved in the phosphotyrosine of target proteins. PP2, a widely used compound to block the activity of Src family kinases, was first used to determine the effect of Src family kinases on TRPM8 phosphotyrosine. A 24 hr exposure of transfected HeLa cells to different concentrations of PP2 (0, 2.5, 10, 20 μM) resulted in a dose-dependent reduction in TRPM8 phosphotyrosine (**Fig. *EV*2, *A* and *B***). In contrast, the reduction was countered by co-application of 1 mM orthovanadate, a protein tyrosine phosphatase inhibitor (**Fig. *EV*2, *C* and *D***). Notably, orthovanadate itself significantly enhanced the basal phosphotyrosine of TRPM8 (**Fig. *EV*2, *C* and *D***). Moreover, we confirmed that only LCK overexpressing, but not BLK and LYN, resulted in a significant increase of TRPM8 phosphotyrosine by about 2.83-fold (**Fig. *EV*2, *E* and *F***), which is consistent the strongest interaction between LCK with TRPM8 (**Fig. 1, *A-C***). These data suggested LCK, as one of Src family kinases, is a potent positive regulator in the regulation of TRPM8 phosphotyrosine via LCK-TRPM8 interaction.

To further provide the document, we performed Co-IP assay to detect the endogenous interaction of TRPM8 and LCK in native cells. Results showed that LCK effectively co-precipitated with TRPM8 in PANC-1 cells (**Fig. 1*D***). Additionally, TRPM8 colocalized with LCK in the cytoplasm by confocal microscopy (**Fig. 1*E***) and purified His-LCK successfully pulled down purified GST-M8C, but not GST alone expressing in BL21 bacteria by protein-protein interaction assay *in vitro* (**Fig. 1*F***). Moreover, cells expressing with increasing amounts of LCK markedly enhanced TRPM8 phosphotyrosine in a dose-dependent manner (**Fig. *EV*2, *G* and *H***) and LCK knockdown significantly decreased TRPM8 phosphotyrosine to about 37% (**Fig. *EV*2, *I* and *J***). Meanwhile, LCK overexpressing markedly enhanced the phosphorylation of C-terminus in TRPM8 by about 2.87-fold in HEK293T cells (**Fig. *EV*2, *K* and *L***) and the immunoprecipitated LCK from transfected HEK293T cells effectively phosphorylated M8C proteins purified from BL21 bacteria by kinase assay *in vitro* (**Fig. *EV*2*M***). We also determine which domain(s) of TRPM8 in the cytoplasm that binds with LCK. Reciprocal Co-IP assays showed that LCK binds with more than one domain of TRPM8 cytosolic (**Fig. 1, *G* and *H***). Together, these data strongly suggested that LCK binds to TRPM8 and positively regulates TRPM8 phosphotyrosine.

### LCK enlarges TRPM8 channel activities

We next employed patch-clamp electrophysiology in HEK293 cells recording whole-cell TRPM8-mediated cation currents (I_TRPMS_) (**Fig. 2*A***), to characterize the functional role of BLK, LCK and LYN in the TRPM8 channel. Compared with the control, overexpressed LCK increased ITRPM8 densities across at depolarization and markedly enlarged ITRPM8 densities by 1.8-fold at +80 mV, whereas BLK and LYN rarely affected (**Fig. 2, *B* and *C***), which was consistent with the above result that only LCK markedly enhanced TRPM8 phosphorylation (**Fig. *EV*2, *E* and *F***). LCK knockdown reduced ITRPM8 densities across at depolarization and markedly decreased I_TRPM8_ densities at +80 mV to about 46% (**Fig. 2, *D* and *E***).

**Figure 2.**
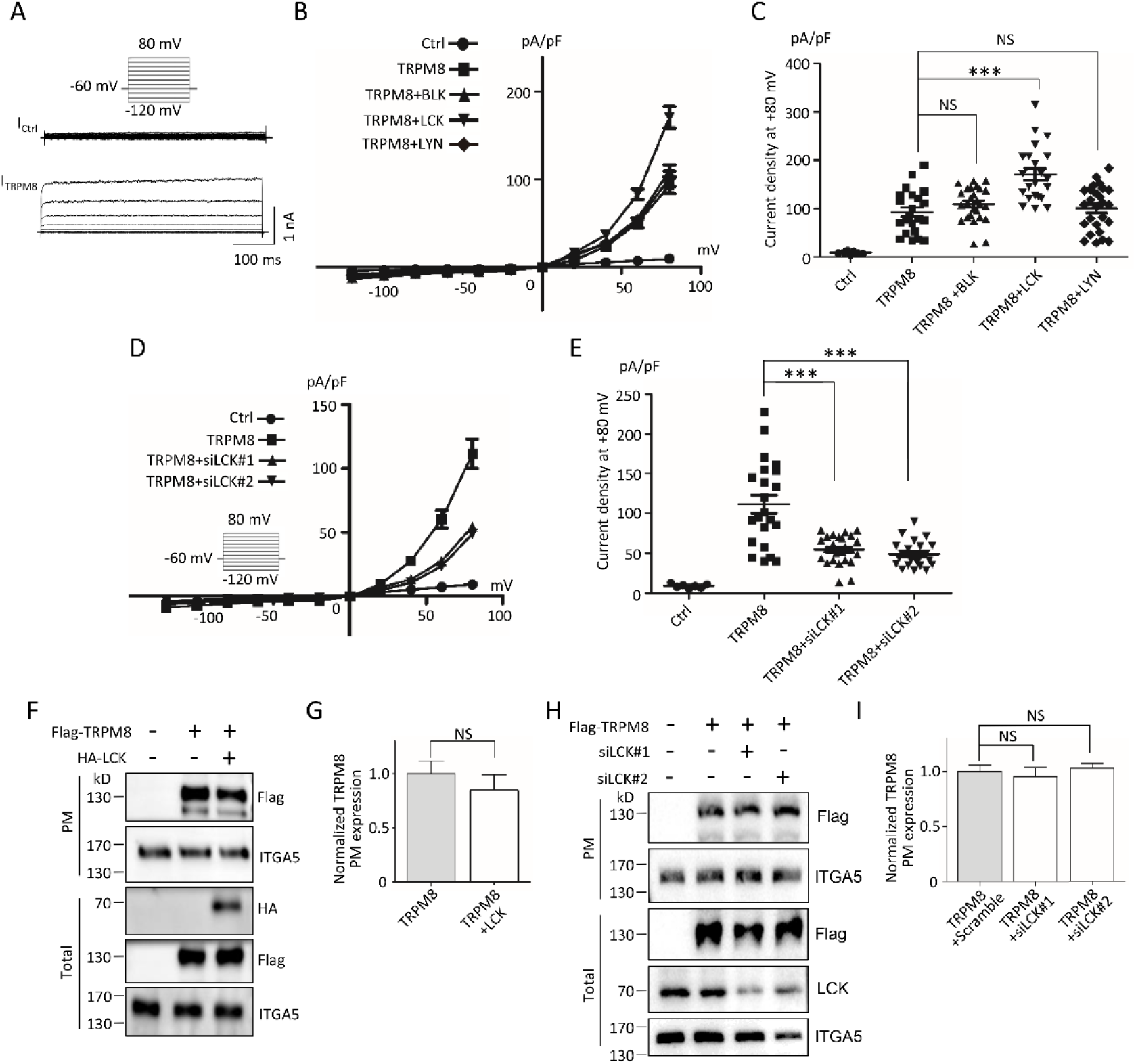
Functional effects of LCK on TRPM8 in HEK293T cells. (**A-E**) Electrophysiological analysis of LCK on whole-cell TRPM8 currents (I_TRPM_s) by patch-clamping experiments. (**A-C**) Expression constructs for Flag-TRPM8 and pEGFP-N1 were co-transfected with HA-BLK, HA-LCK, HA-LYN, or control vector into HEK293T cells. EGFP-positive cells were selected for recording I_TRPM8_ (n=5~20 cells per group). The voltage clamp protocol is shown in the *inset* of figure. (**A**) Representative imaging of I_TRPM8_. (**B**) The relationship of average I_TRPM8_ density (I_TRPM8_ normalized to cell capacitance) and voltage. (**C**) Quantification of peak I_TRPM8_ density on +80 mV as in ***B***. (**D-E**) Similar I_TRPM8_ recordings as in ***A*** but HEK293T cells co-expressing Flag-TRPM8 and pEGFP-N1 with siLCK or negative scramble siRNAs. (n=5~20 cells per group). (**D**) The relationship of average I_TRPM8_ density and voltage. (**E**) Quantification of peak I_TRPM8_ density on +80 mV as in ***D***. (**F-I**) Cell-surface biotinylation assays for detecting TRPM8 PM expression. (**F**) Representative WB images of TRPM8 on the PM and total lysates from HEK293T cells co-transfected with or without HA-LCK. (**G**) Quantification of PM and total protein expression levels of TRPM8 in *(**F***). (**H-I**) Similar experiments in ***F*** and **G** but cells co-expressing with siLCK or negative scramble siRNAs. ITGA5 (Intergrin α5) was used as a loading control for PM proteins. ***, P < 0.001, NS, not significant. Data are presented as mean ± SEM. All studies were repeated at least three times.

To assess the molecular mechanism for the enlarged ITRPM8 by LCK through enhancing TRPM8 expression on PM, we extracted TRPM8 proteins on the PM from transfected HEK293T cells. Results showed that overexpressed LCK as well as LCK knockdown rarely affected the PM and total expression of TRPM8 amount (**Fig. 2, *F-I***), suggesting that LCK modulated TRPM8 channel function likely through regulating the biophysical properties of TRPM8 channel, but not PM TRPM8 trafficking. These data together indicated that LCK acts as a positive regulator of TRPM8-mediated currents.

### LCK affects the multimerization but not intramolecular N-C binding of TRPM8

Due to LCK interaction with both N terminus and C terminus of TRPM8 (**Fig. 1, *G* and *H***) and the importance of the intramolecular N-C binding for activation of TRPM8 function (Rohacs et al, 2005; Zheng et al, 2018), we investigated whether LCK affected the intramolecular N-C binding of TRPM8. Results showed that overexpressed LCK rarely affected the intramolecular N-C binding of TRPM8 with or without PP2 (**Fig. 3, *A* and *B***). Moreover, the intramolecular N-C binding of TRPM8 was not altered in the presence of LCK knockdown (**Fig. 3, *C* and *D***). We next investigated whether LCK modulated TRPM8 multimerization, which was critical for TRPM8 channel function as previously reported (Erler et al, 2006; Stewart et al, 2010). Results showed that overexpressed LCK markedly enhanced, whereas LCK knockdown decreased TRPM8 multimerization in the presence of DSS (**Fig. 3, *E-H***). To provide further document, we also detected the effect of LCK on the binding of intermolecular TRPM8. Results showed that LCK overexpression markedly increased the binding of intermolecular TRPM8, while the increase was significantly reduced by co-application of PP2 (**Fig. 3, *I* and *J***). Moreover, LCK knockdown effectively reduced the binding of intermolecular TRPM8 (**Fig. 3, *K* and *L***). These data together indicated that LCK affects TRPM8 multimerization but not intramolecular N-C binding, thereby regulating its channel functions.

**Figure 3.**
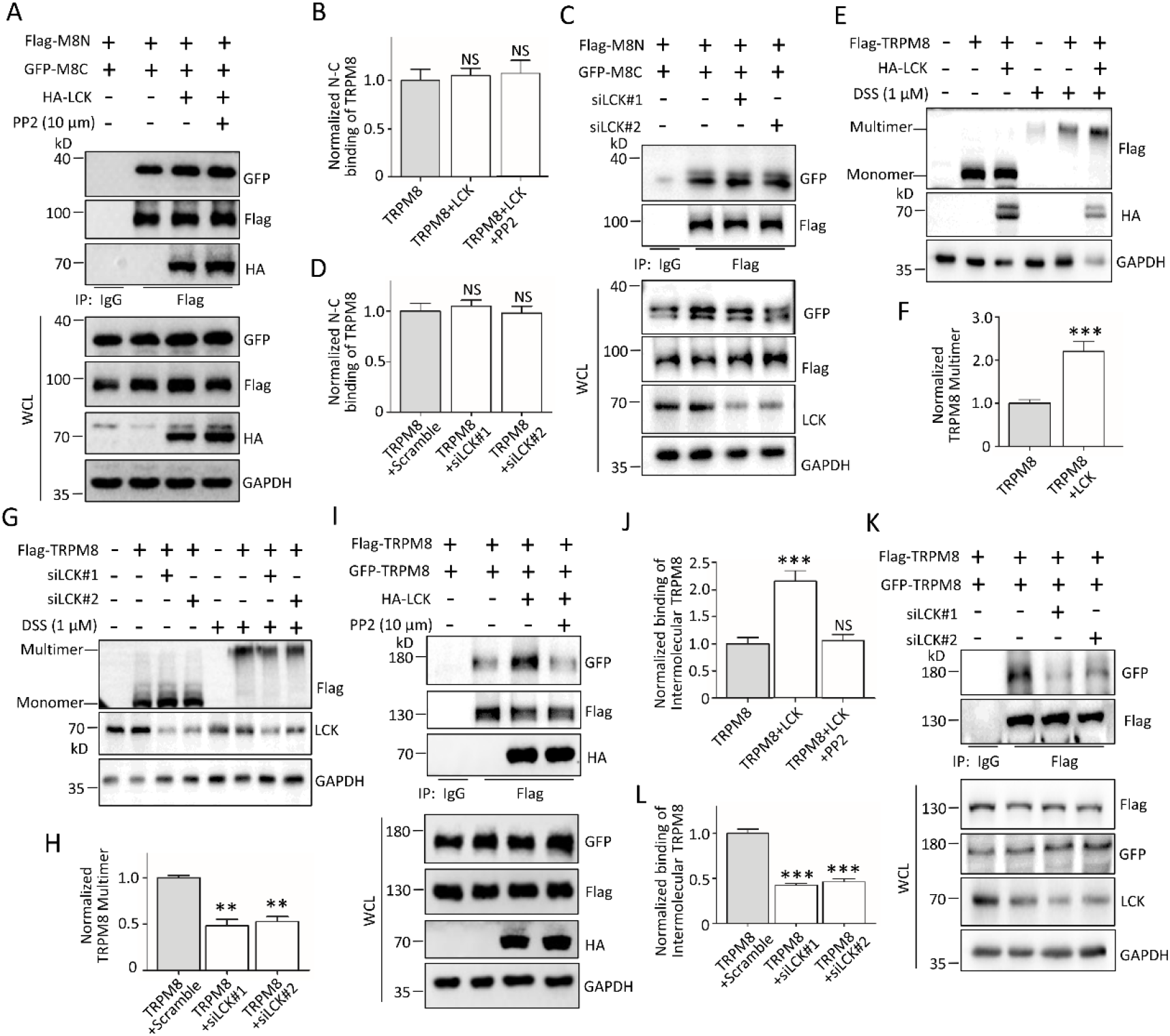
The multimerization, but not intracellular N-C binding of TRPM8, regulated by LCK. (**A-D**) The effect of LCK on the intracellular N-C binding of TRPM8. (**A-B**) Expression constructs for Flag-tagged N-terminus of TRPM8 (Flag-M8N) and GFP-M8C were co-transfected with HA-LCK or control vector into HEK293T cells, before harvest for treatment with 10 μM PP2 for 24 h. The cells were then harvested for IP with an anti-Flag antibody and WB assay with the indicated antibodies to determine the intracellular N-C binding of TRPM8. (**C-D**) Similar experiments in ***A*** and ***B*** but cells co-expressing Flag-M8N and GFP-M8C along with siLCK. (**E-L**) The effect of LCK on the multimerization of TRPM8. (**E-F**) Expression constructs for Flag-TRPM8 were co-transfected with HA-LCK or control vector into PANC-1 cells, before harvest for treatment with 1 μM DSS for 30 min, a crosslinking agent. The cell lysates were subjected to WB assay with the indicated antibodies to detect the level of TRPM8 multimerization. (**G-H**) Similar experiments in ***E*** and ***F*** but cells co-expressing Flag-TRPM8 along with siLCK. (**I-J**) Flag-TRPM8 and GFP-TRPM8 were co-transfected with or without HA-LCK into AsPC-1 cells. The cells were then harvested for IP with an anti-Flag antibody and WB assay with the indicated antibodies to determine the binding of intermolecular TRPM8. (**K-L**) Similar experiments in ***I*** and ***J*** but cells co-expressing Flag-TRPM8 and GFP-TRPM8 along with siLCK. **, P < 0.01, ***, P < 0.001, NS, not significant. Data are presented as mean ± SEM. All studies were repeated at least three times.

### LCK enhances phosphotyrosine of TRPM8 at Y1022

We next determined which of the potential phosphotyrosine site(s) in TRPM8 regulated by LCK. Using the combination of immunoprecipitation and MS analysis, lysine residue at 423 was identified to be a major ubiquitination site of TRPM8 earlier reported by us (Huang et al, 2021). Within the same screen, the highly conserved tyrosine residue at 1022 across species was also detected to be a potential phosphotyrosine site in TRPM8 (**Fig. 4, *A* and *B***). In order to confirm and characterize the importance of Y1022 for TRPM8 phosphotyrosine, the expression construct for single point mutant TRPM8-Y1022F was generated. Compared to wild type TRPM8 (WT-TRPM8), mutant TRPM8-Y1022F significantly reduced its phosphotyrosine level with or without LCK overexpression (**Fig. 4, *C* and *D***). The significant differences of phosphorylation level between WT-TRPM8 and mutant TRPM8-Y1022F in the presence of LCK overexpression were abolished by saracatinib (**Fig. 4, *C* and *D***), a potent and selective inhibitor of Src-family tyrosine kinases (SRC, YES and LCK) (Hennequin et al, 2006). Kinase assay *in vitro* (**Fig. 4, *E* and *F***) further supported the conclusion that Y1022 in TRPM8 is the potent phosphorylation target for LCK. Interestingly, we found that mutant TRPM8-Y1022F significantly decreased own serine phosphorylation (phosphoserine), but not altering threonine phosphorylation (phosphothreonine) of TRPM8 (**Fig. *EV*3, *A* and *B***). Moreover, LCK significantly enhanced TRPM8 phosphoserine and phosphotyrosine, but not phosphothreonine (**Fig. *EV3, C* and *D***).

**Figure 4.**
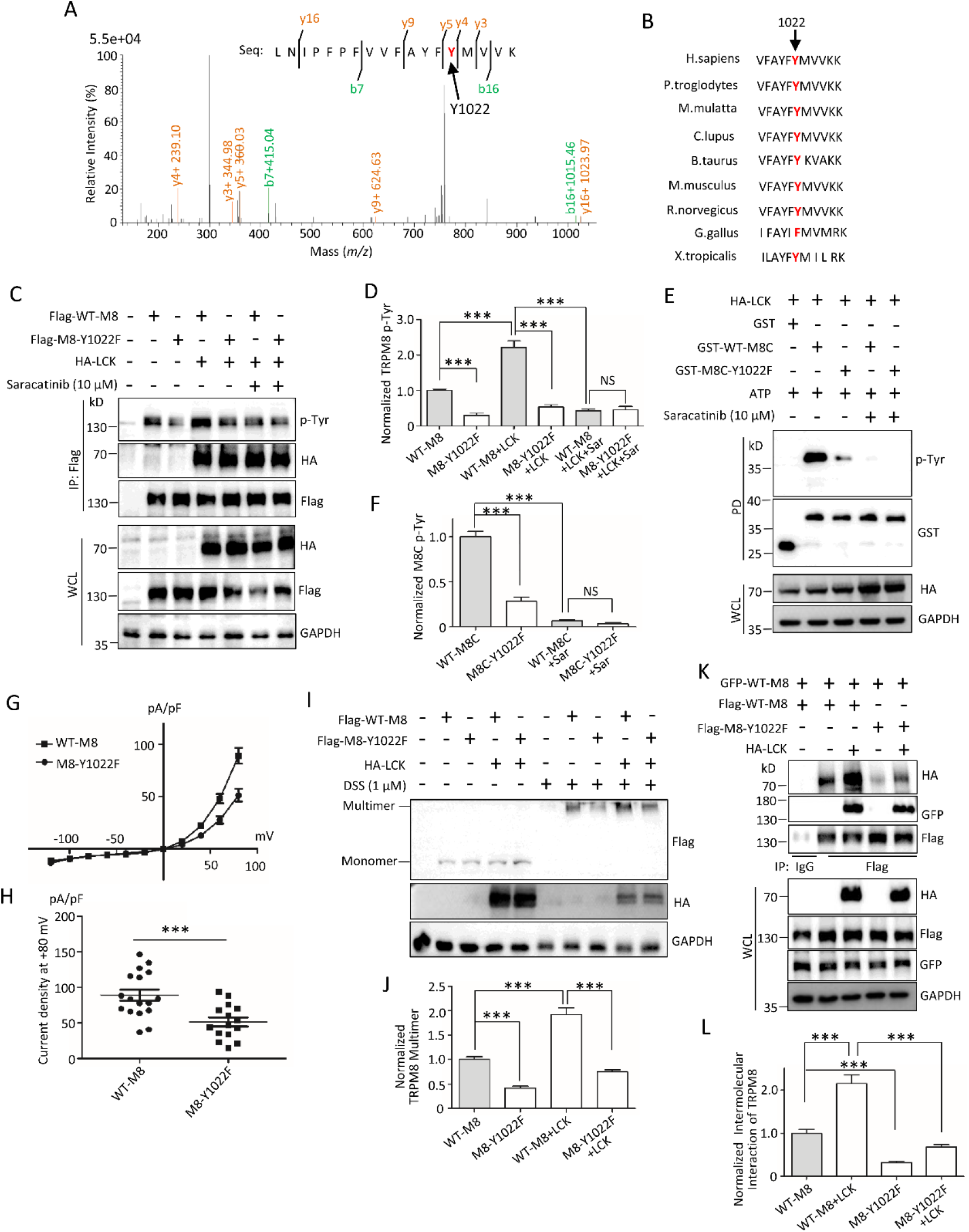
Identification of phosphotyrosine on TRPM8 at position 1022 regulated by LCK. (**A**) MS imaging of phosphotyrosine site of TRPM8 in combination with the NCBI blast (peptide sequences are indicated). (**B**) Amino acid sequence alignment showing that tyrosine at position 1022 is highly conserved among multiple species. (**C-D**) Expression constructs for Flag-tagged wild type TRPM8 (Flag-WT-M8) or mutant Y1022F (Flag-M8-Y1022F) were transfected with or without HA-LCK into HEK293T cells, before harvest for treatment with 10 μM saracatinib for 24 h. The lysates were then used for IP with an anti-Flag antibody and then subjected to WB assay with the indicated antibodies to detect the level of TRPM8 phosphotyrosine. (**E-F**) Kinase assay *in vitro.* Purified GST alone, GST tagged wild type or mutant of C-terminus of TRPM8 fusion proteins expressing in *E.coli* bacteria were mixed with HA-LCK immunoprecipitated with anti-HA antibody from HEK293T cells expressing HA-LCK construct, 1 mM ATP or their combination in kinase assay buffer to determine the level of M8C phosphotyrosine. (**G**) Relationship of test potential and averaged densities of I_TRP_M8 recorded from HEK293T cells co-transfected Flag-WT-M8 or Flag-M8-Y1022F with pEGFP-N1. (**H**) Peak current density on +80 mV as in ***G*** (n=15~20 cells per group). (**I-J**) Expression constructs for Flag-WT-M8 or Flag-M8-Y1022F were co-transfected with or without HA-LCK into PANC-1 cells, before harvest for treatment with 1 μM DSS for 30 min for WB to detect the level of TRPM8 multimerization. (**K-L**) Flag-WT-M8 or Flag-M8-Y1022F along with GFP-tagged wild type TRPM8 (GFP-WT-M8) were co-transfected with or without HA-LCK into AsPC-1 cells to determine the binding of intermolecular TRPM8. ***, P < 0.001, NS, not significant. Data are presented as mean ± SEM. All studies were repeated at least three times.

We next recorded the cation currents mediated by mutant TRPM8-Y1022F in HEK293T cells and found mutant TRPM8-Y1022F decreased I_TRPM8_ densities across at depolarization when compared with WT-TRPM8 (**Fig. 4*G***). At +80 mV, mutant TRPM8-Y1022F markedly decreased ITRPM8 densities to about 53% (**Fig. 4*H***). In addition, mutant TRPM8-Y1022F did not affect the expression of total and PM TRPM8 with or without LCK (**Fig. *EV*3, *E-G***). WB assay showed that mutant TRPM8-Y1022F significantly reduced TRPM8 multimerization with or without LCK when compared to WT-TRPM8 (**Fig. 4, *I* and *J***). The mutant TRPM8-Y1022F significantly impaired the interaction of intermolecular TRPM8 (**Fig. 4, *K* and *L***), further supporting that the importance of Y1022 for TRPM8 multimerization. Together, these data suggested mutant TRPM8-Y1022F modulated channel function likely through modulating TRPM8 multimerization, which are consistent with the above findings that the effects of LCK on TRPM8 function and multimerization.

### 14-3-3ζ mediates LCK in the regulation of TRPM8 multimerization

The 14-3-3 protein is a widely expressed acidic protein that binds with phosphorylated targeted proteins and enhances its multimerization for regulating its activities (Clatot et al, 2017). 14-3-3ζ, a member of the 14-3-3 protein family, was identified in the same screen at ~30 kD bands as our previous studies in Fig. 1a (Huang et al, 2021) (**Fig. *EV*1, *E* and *F***). We first demonstrated the interaction of 14-3-3ζ and the C terminus of TRPM8 by an *in vitro* GST pull-down assay (**Fig. 5*A***) and 14-3-3ζ and TRPM8 in native PANC-1 cells by Co-IP assay (**Fig. 5*B***), suggesting that TRPM8 interacted with 14-3-3ζ in a protein complex. Next, we determined the functional role of 14-3-3ζ on TRPM8 multimerization. Results showed that 14-3-3ζ overexpressing or knockdown, respectively, markedly enhanced or reduced TRPM8 multimerization in the presence of DSS (**Fig. 5, *C-F***). 14-3-3ζ knockdown effectively inhibited the binding of intermolecular TRPM8 in AsPC-1 cells (**Fig. 5, *G* and *H***), further supporting the role of 14-3-3ζ on TRPM8 multimerization.

**Figure 5.**
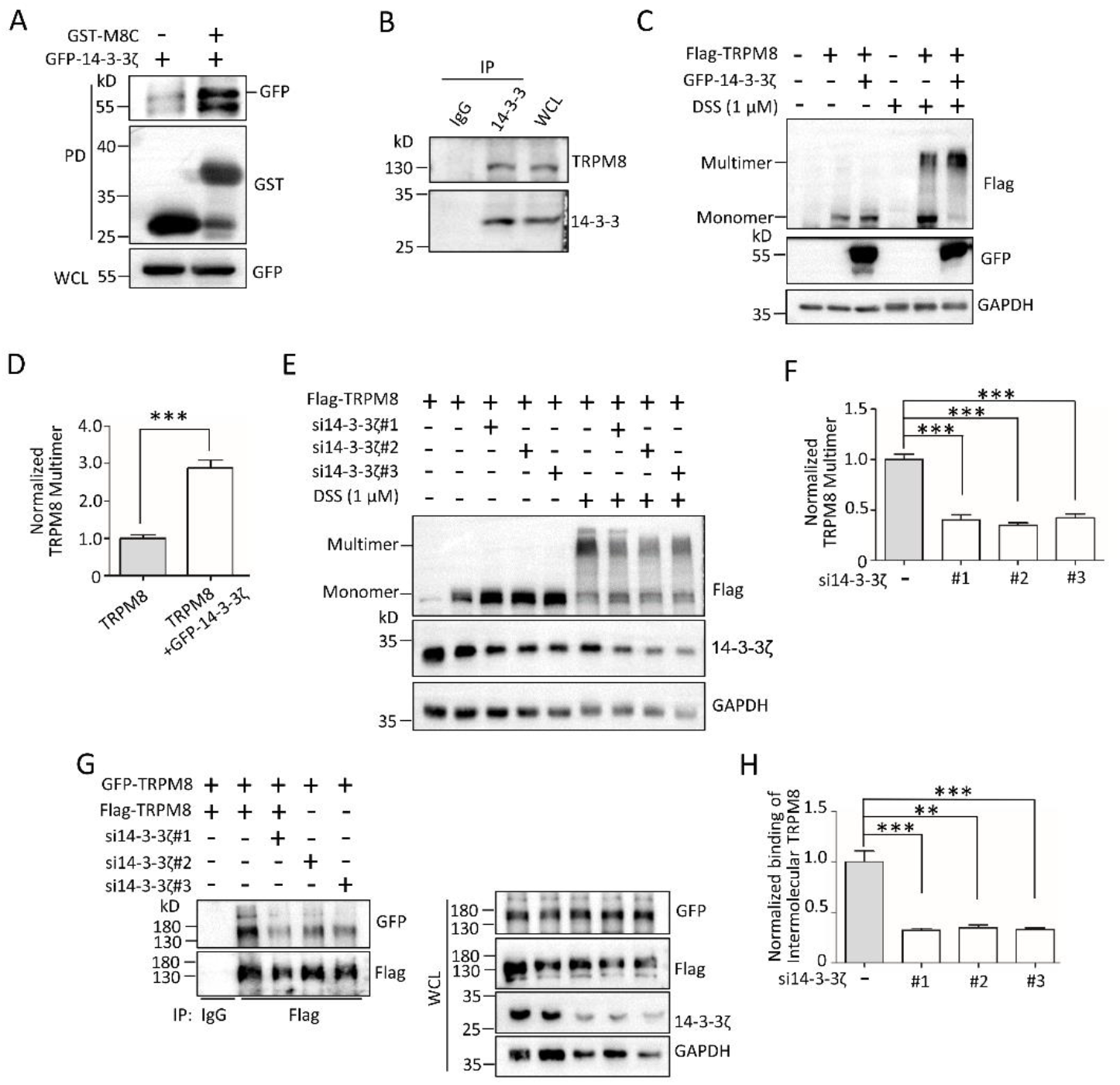
14−3−3ζ interaction with TRPM8 for regulating TRPM8 multimerization. (**A**) GST pull-down assay. Purified GST alone or GST-M8C fusion proteins expressing in *E.coli* BL21 bacteria were incubated with the lysates from HEK293T cells expressing GFP-14−3−3ζ constructs and subjected to WB assay. (**B**) Co-IP assay. The lysates of native PANC-1 cells were added to an anti-14−3−3 antibody for IP and then subjected to WB assay with an anti-TRPM8 antibody. (**C-D**) Expression constructs for Flag-TRPM8 with or without GFP-14−3−3ζ were transfected into PANC-1 cells, before harvest for treatment with 1 μM DSS for 30 min. The lysates were subjected to WB assay with the indicated antibodies to detect the level of TRPM8 multimerization. (**E-F**) Similar experiments in ***C*** and ***D*** but cells co-expressing Flag-TRPM8 along with human 14−3−3ζ-specific siRNAs (si14−3−3ζ#1, #2, or #3). (**G-H**) AsPC-1 cells were co-transfected Flag-TRPM8 and GFP-TRPM8 with or without si14−3−3ζ, and then harvested for IP with an anti-Flag antibody and WB assay with the indicated antibodies to determine the binding of intermolecular TRPM8. **, P < 0.01, ***, P < 0.001. Data are presented as mean ± SEM. All studies were repeated at least three times.

Based on the above findings, we hypothesized that 14-3-3ζ involved in the regulation of LCK on TRPM8 multimerization. We first determined whether LCK-mediated TRPM8 phosphotyrosine affected the binding of TRPM8 and 14-3-3ζ. Results showed that LCK overexpression and knockdown, respectively, significantly increased and reduced the binding of TRPM8 and 14-3-3ζ (**Fig. 6, *A-D***). Meanwhile, mutant TRPM8-Y1022F significantly impaired the binding of 14-3-3ζ and TRPM8 in the presence or absence of LCK (**Fig. 6, *E* and *F***). These data suggested that TRPM8 phosphotyrosine regulated by LCK positively modulated 14-3-3ζ-TRPM8 interaction. Moreover, 14-3-3ζ knockdown eliminated the function of LCK on an increase of TRPM8 multimerization (**Fig. 6, *G* and *H***), which is further validated by the impaired interaction of intermolecular TRPM8 regulated by LCK in the presence of 14-3-3ζ knockdown (**Fig. 6, *I* and *J***). Together, these data revealed that 14-3-3ζ is critical for LCK on the regulation of TRPM8 multimerization.

**Figure 6.**
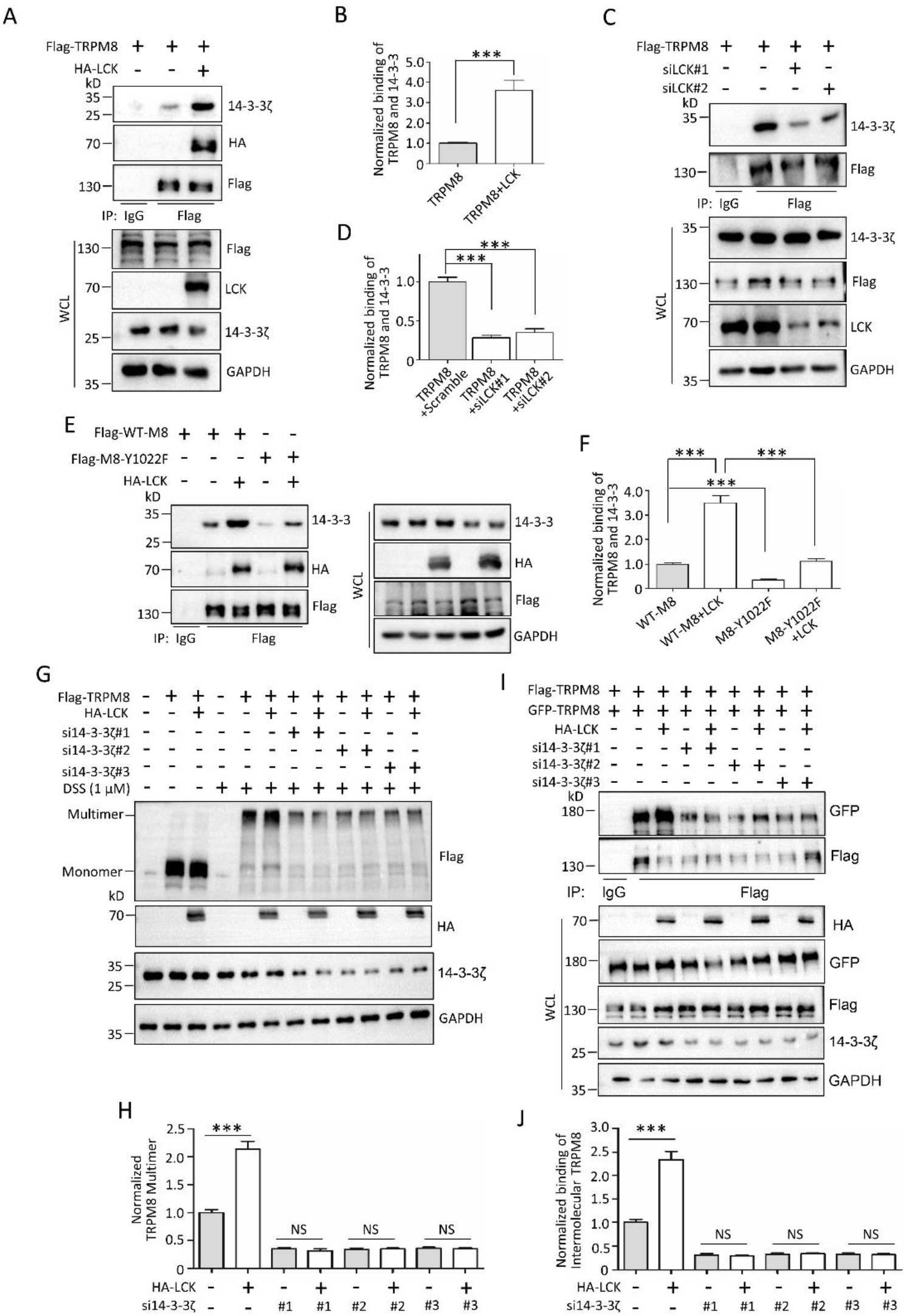
14−3−3ζ involved in the regulation of LCK on the multimerization of TRPM8. (**A-B**) PANC-1 cells were co-transfected Flag-TRPM8 with HA-LCK or control vector and then harvested for IP with an anti-Flag antibody and WB assay with the indicated antibodies to determine the binding of TRPM8 and 14−3−3ζ. (**C-D**) Similar experiments in ***A*** and ***B*** but cells co-expressing Flag-TRPM8 along with siLCK. (**E-F**) Expression constructs for Flag-WT-TRPM8 or Flag-TRPM8-Y1022F along with GFP-TRPM8 were co-transfected with or without HA-LCK into HEK293T cells to determine the binding of intermolecular TRPM8. (**G-H**) PANC-1 cells were co-transfected with Flag-TRPM8, HA-LCK, si14−3−3ζ (#1, #2 or #3) or their combination, before harvest for treatment with 1 μM DSS for 30 min, and subjected to WB assay to detect the level of TRPM8 multimerization. (**I-J**) AsPC-1 cells were co-transfected with Flag-TRPM8 and GFP-TRPM8, HA-LCK, si14−3−3ζ or their combination, and then harvested for IP with an anti-Flag antibody and subjected to determine the binding of intermolecular TRPM8. ***, P < 0.001, NS, not significant. Data are presented as mean ± SEM. All studies were repeated at least three times.

### TRPM8 phosphotyrosine positively modulates LCK activity

Phosphorylation on Tyr394 or Tyr505 is critical for regulation of LCK activity (Eck et al, 1994; Peri et al, 1993; Yamaguchi & Hendrickson, 1996), which was also validated by our data that the mutant Y394F and Y505F, respectively, significantly inhibited and enhanced the function of LCK on TRPM8 phosphotyrosine (**Fig. *EV*4, *A* and *B***). Next, we detected whether TRPM8 phosphotyrosine feedback modulated LCK activity, phosphorylation-site-specific antibodies of LCK on Y394 and Y505 was used to detect the level of LCK Y394 and Y505 phosphorylation, which were verified to be specific and effective (**Fig. *EV*5 *A***). Results revealed that the level of phosphorylated LCK on Y394 was comparable between cells overexpressing control vector, WT-TRPM8 and TRPM8-Y1022F (**Fig. 7, *A* and *B***). However, WT-TRPM8 markedly reduced the level of phosphorylated LCK on Y505 compared to control, while the reduction was countered in the presence of mutant Y1022F-TRPM8 (**Fig. 7, *A* and *B***). Meanwhile, TRPM8 rarely affected LCK Ser/Thr phosphorylation (**Fig. *EV*5, *B* and *C***). These data together suggested that TRPM8 phosphotyrosine positively modulated LCK activity through inhibition of phosphorylated LCK on Y505. Meanwhile, TRPM8 overexpressing significantly decreased the level of phosphorylated LCK on Y505 but not Y394, with or without saracatinib. However, saracatinib markedly enhanced the level of phosphorylated LCK on Y505 (**Fig. *EV*5, *D* and *E***), speculating that saracatinib inhibited LCK activity likely through activation of Tyr505 phosphorylation.

**Figure 7.**
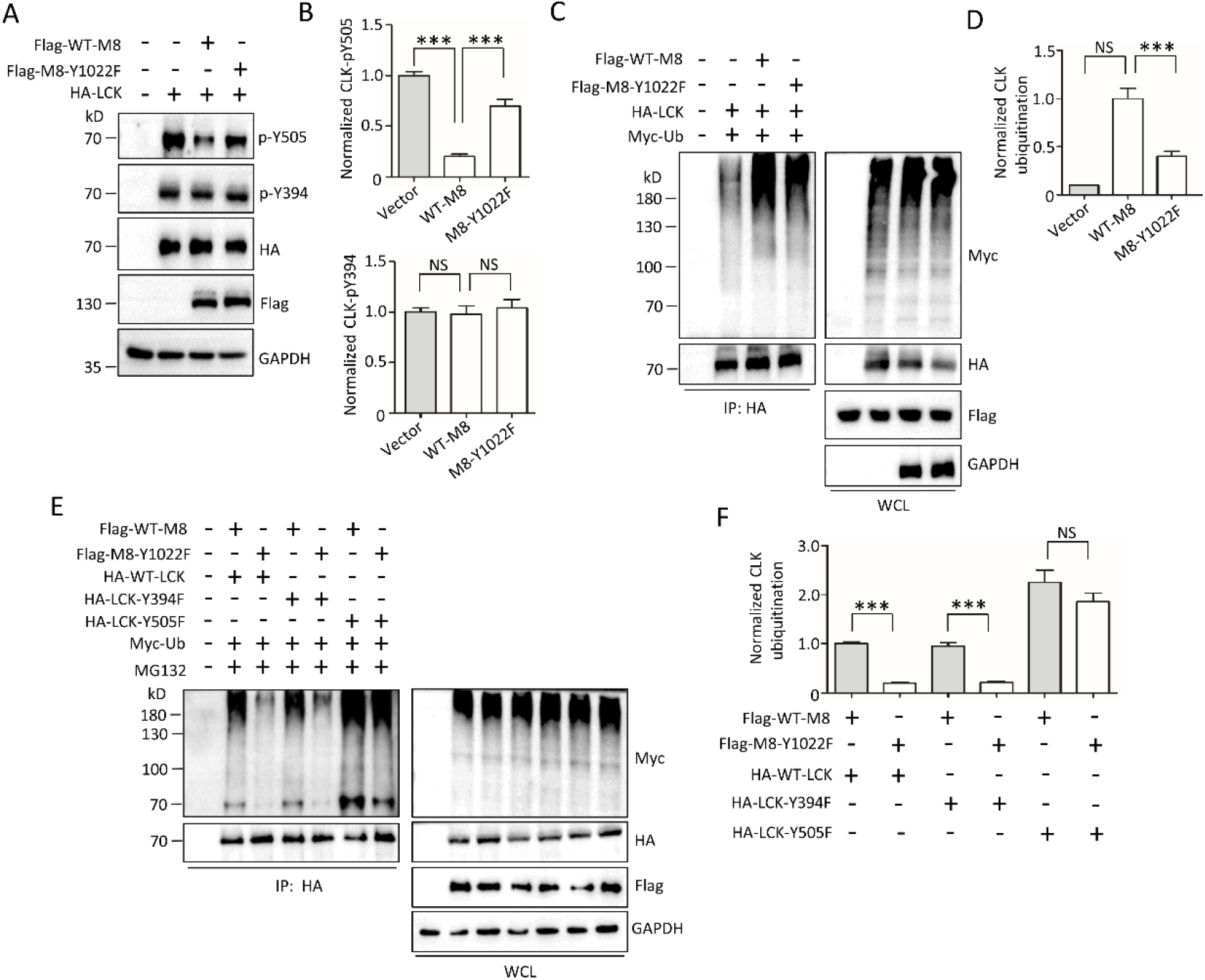
LCK activity regulated by TRPM8 phosphotyrosine. (**A-B**) LCK phosphorylation assay. PANC-1 cells were co-transfected HA-LCK with control vector, WT-TRPM8 or mutant TRPM8-Y1022 with the indicated antibodies as shown in (**A)**, and quantification of the level of LCK phosphorylation shown in (**B**). (**C-F**) LCK ubiquitination assay. (**C-D**) Expression constructs for HA-LCK and Myc-Ub were co-transfected with control vector, WT-TRPM8 or mutant TRPM8-Y1022 into AsPC-1 cells. The cells were treated with 10 μM MG132 for 6 h before harvest and used for IP with an anti-HA antibody and WB with the indicated antibodies. (**E-F**) Expression constructs for Myc-Ub and WT-TRPM8 or mutant TRPM8-Y1022F were co-transfected with HA-tagged wild type or mutant LCK into PANC-1 cells. The cells were treated with 10 μM MG132 before harvest and used for IP with an anti-HA antibody and WB with the indicated antibodies. ***, P < 0.001, NS, not significant. Data are presented as mean ± SEM. All studies were repeated at least three times.

In addition to phosphorylation, ubiquitination is also involved in the regulation of LCK activity (Eck et al, 1994; Peri et al, 1993; Yamaguchi & Hendrickson, 1996). Ubiquitination assay showed that WT-TRPM8 overexpression significantly increased LCK ubiquitination than control vector, whereas the increase was partially impaired in the presence of mutant Y1022F-TRPM8 TRPM8 (**Fig. 7, *C* and *D***), suggesting that TRPM8 phosphotyrosine feedback modulated LCK ubiquitination. Moreover, we also detected the effect of TRPM8 phosphotyrosine on the ubiquitination of LCK mutants Y394F and Y505F. Compared to WT-TRPM8, mutant TRPM8-Y1022F showed similar inhibitory effects on the ubiquitination of WT-LCK and mutant LCK-Y394F (**Fig. 7, *E* and *F***). However, the inhibitory effect of TRPM8-Y1022F on LCK ubiquitination was almost abrogated in the presence of mutant LCK-Y505F (**Fig. 7, *E* and *F***), suggesting that TRPM8 phosphorylation modulated the ubiquitination of the inactive form of LCK (LCK-Y394F).

### Impaired phosphorylation of TRPM8 inhibits pancreatic cancer cell proliferation, migration and tumorigenesis *in vitro* and *in vivo*

We next examined the effect of Y1022 in TRPM8 on pancreatic cancer cell proliferation and migration by use of RFP labeled cell lines of PANC-1 or AsPC-1 cells stably expressing control vector, WT-TRPM8, or mutant TRPM8-Y1022F. EdU incorporation assay, immunofluorescence and colony formation together revealed that WT-TRPM8 significantly increased tumor cells proliferation when compared to control vector cells. However, mutant TRPM8-Y1022F impaired the function of TRPM8 on increased cells proliferation (**Fig. 8, *A-D***). These data suggested that TRPM8 phosphorylation at Y1022 is critical for pancreatic cancer cell proliferation.

**Figure 8.**
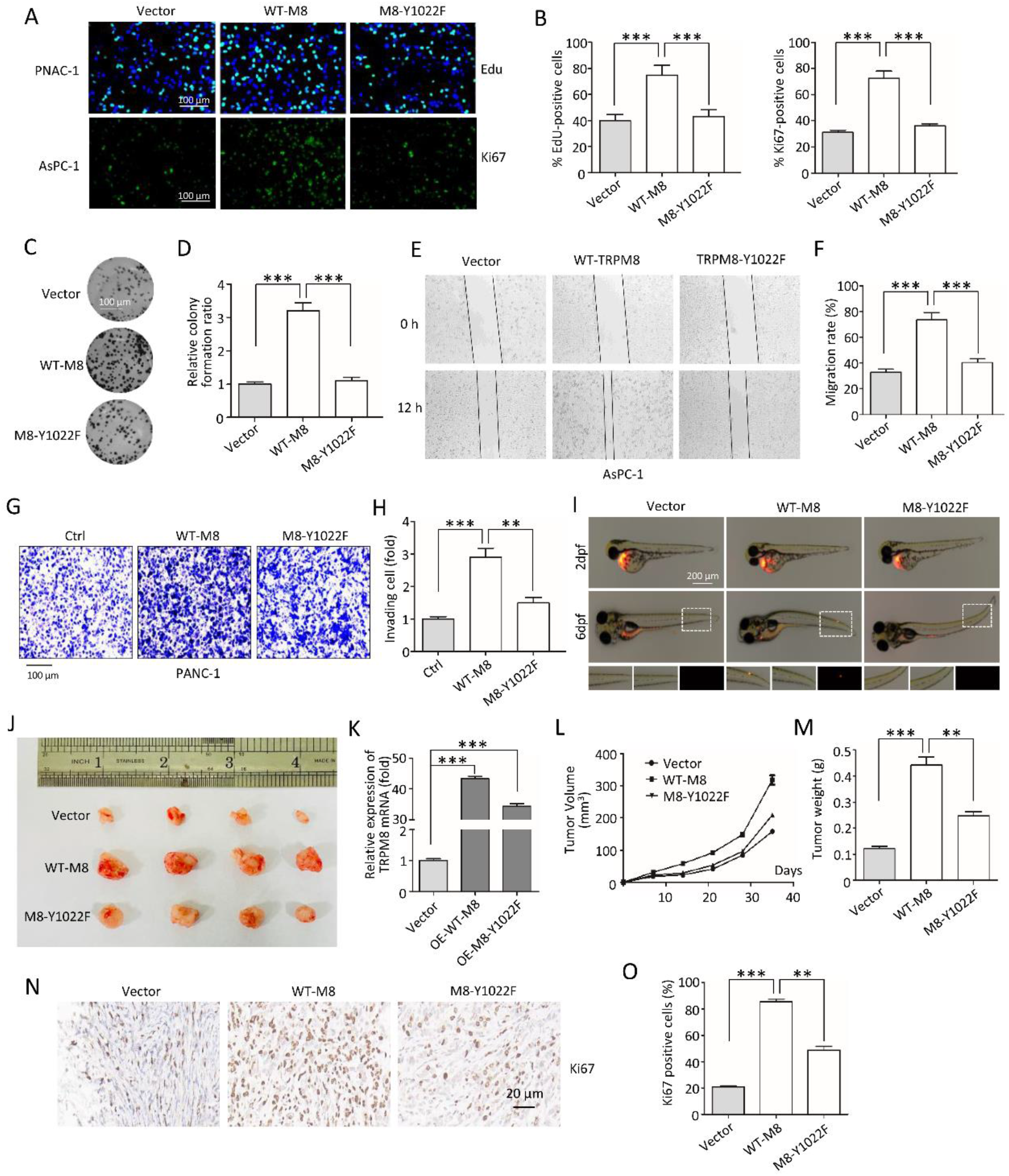
The role of Y1022 on TRPM8 on tumor cell proliferation, migration and tumorigenesis. (**A-D**) Cell proliferation assays *in vitro.* (**A-B**) The RFP labeled cell lines of PANC-1 or AsPC-1 cells stably expressing control vector, WT-TRPM8, or mutant TRPM8-Y1022F were constructed and used for EdU incorporation assays (Upper panel) and Ki67 immunofluorescence (Lower panel). Scale bars, 100 μm. (**C-D**) Colony formation assays were performed in RFP labeled PANC-1 stably maintained cells. Scale bars, 100 μm. (**E-I**) Cell migration assays. (**E-F**) Wound-healing assay was performed in RFP labeled AsPC-1 stably maintained cells. (**G-H**) Transwell assay was performed in RFP labeled PANC-1 stably maintained cells. Scale bars, 100 μm. (**I-L**) Animal xenotransplantation engraftment experiments. (**I**) Representative confocal microscopy images of 6 days xenotransplantation of zebrafish injecting with RFP labeled PANC-1 stably maintained cells. Scale bars, 200 μm. (**J**) Imaging of tumors excised from the mice subcutaneously injecting RFP labeled PANC-1 stably maintained cells by growth for 5 weeks. (**K**) Quantification of the expression of TRPM8 mRNA in (***J***). (**L**) Weights of the excised tumors in each group in (***J***). (**M**) Growth curves showing the changes in the tumor volume in mice in different groups every 5 days from the injection. (**N**) Representative H&E staining images and immunohistochemical images of Ki67 in excised tumors tissues. Scale bars, 20 μm. (**O**) Quantification of Ki67 expression in (***N***). **, P < 0.01, ***, P < 0.001. Data are presented as mean ± SEM. All studies were repeated at least three times.

In addition, wound-healing and transwell assays revealed that WT-TRPM8 showed a significantly higher migration capacity than control vector cells, whereas mutant TRPM8-Y1022F impaired the function of TRPM8 on cell migration (**Fig. 8, *E-H***). We also employed a novel metastatic zebrafish xenotransplantation model to detect mutant TRPM8-Y1022F on tumor cell migration. After 6 days of xenotransplantation, compared to control group, a massive number of PANC-1 cells stably expressing WT-TRPM8 migrated to distant parts of the zebrafish body to form micrometastasis, while PANC-1 cells stably expressing mutant TRPM8-Y1022F did not migrate far from the primary site (**Fig. 8*I***). These data together suggested that TRPM8 phosphorylation at Y1022 is critical for pancreatic cancer cell migration.

**Figure 9.**
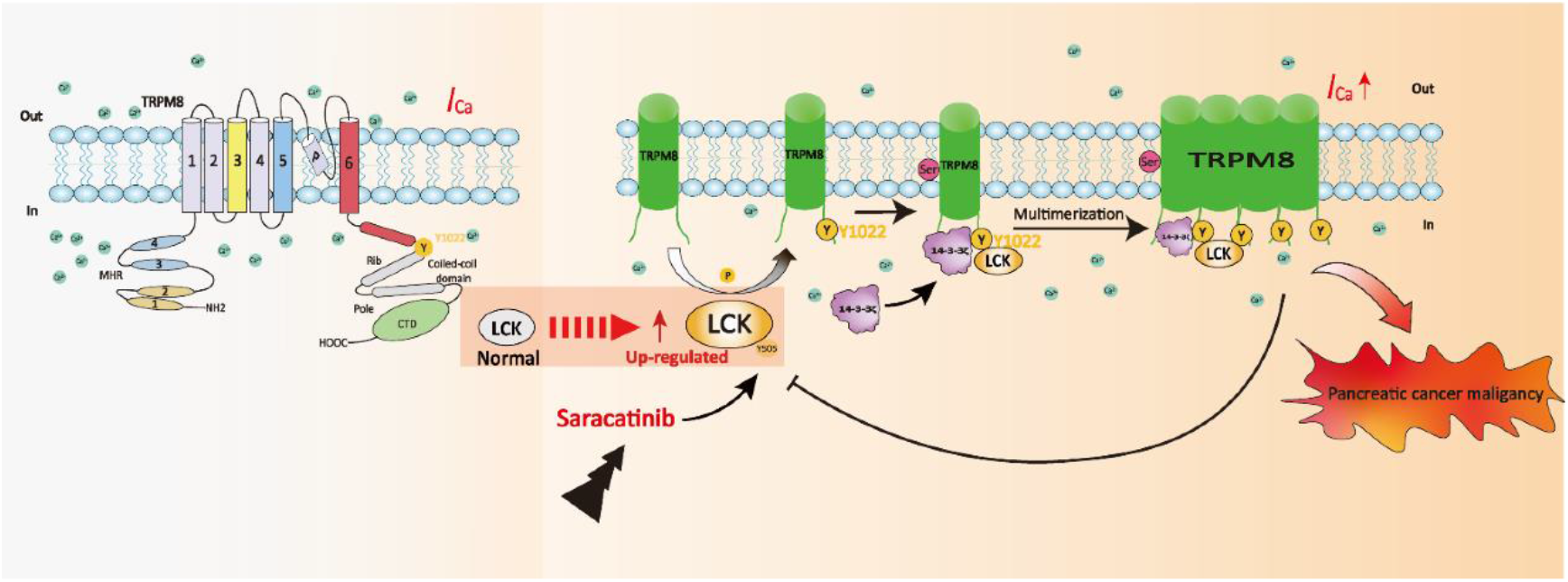
Schematic diagram of the biological role of the LCK-14−3−3ζ-TRPM8 axis in egulation of TRPM8 function and LCK activity.

To further assess the importance of TRPM8 phosphorylation at Y1022 tumorigenesis *in vivo,* BALB/c nude mice bearing subcutaneous pancreatic xenograft tumors derived from PANC-1 cells stably expressing control vector, WT-TRPM8, and mutant TRPM8-Y1022F. After 35 days of growth, the tumors were carefully taken out (**Fig. 8*J***). TRPM8 mRNA expression detected by real time qRT-PCR was up-regulated in the tumor xenografts that injecting cells with stable expression of WT-TRPM8 or TRPM8-Y1022F (**Fig. 8*K***). Moreover, compared to the control xenograft tumors, a significant increase on tumor volumes, weights were observed in WT-TRPM8 xenograft tumors and MKI67 expression was markedly increased in WT-TRPM8 xenograft tumor tissues by histopathologic analyses (**Fig. 7, *L-O***). However, mutant TRPM8-Y1022F diminished the increase of WT-TRPM8 on tumor volumes and weights as well as MKI67 expression. These data together suggested that TRPM8 phosphorylation at Y1022 contributes to tumorigenesis *in vivo*.

## Discussion

TRPM8, which function as a Ca^2+^-permeable channel, is required to assembly to form functional homologous tetramers (Erler et al, 2006; Phelps & Gaudet, 2007; Tsuruda et al, 2006) and plays a vital role in environmental cold sensing, menthol-induced analgesia of acute and inflammatory pain, and migraine (Liu et al, 2013). Elevated expression of TRPM8 has been found in human pancreatic cancer and several other diseases in clinical patients (Liu et al, 2016). However, how TRPM8 channel in PM exercises its oncogenic effects are not well understood. Moreover, phosphotyrosine is involved in the regulation of TRPM8 function without identification of the exact site(s) (Manolache et al, 2020). In the present study, we identified LCK and 14-3-3ζ as new TRPM8 binding partners and a novel post-translational modification of TRPM8 at Y1022. Moreover, we provided a novel model that LCK mediated TRPM8 phosphotyrosine at Y1022 is critical for ITRPM8 densities through enhancing 14-3-3ζ-TRPM8 binding to regulate TRPM8 multimerization. Knockdown of 14-3-3ζ markedly impaired the regulation of LCK on TRPM8 multimerization. Additionally, phosphorylation and ubiquitination mediated LCK activity was coordinately regulated by TRPM8 phosphotyrosine at Y1022 in a feedback loop. Importantly, we provided multiple lines of evidence supporting the importance of TRPM8 phosphotyrosine at Y1022 on pancreatic cancer progression *in vitro* and *in vivo*.

Earlier studies showed that the member of Src family kinases Src, but not Abl and Btk, phosphorylates TRPM8 and modulates the cold-induced activation of TRPM8 channel by using a combination of a constitutively active isoform of Src, Src inhibitor, and Src siRNA, without detecting the interaction of TRPM8 and Src (Manolache et al, 2020). In this study, we employed GST pull-down in combination with MS assays and found BLK, LCK, and LYN were potential interacting partners of TRPM8, which is further strengthened by Co-IP assays. However, ~60 kD of Src was failed to detect in a similar size band of BLK, LCK, and LYN. We speculated that Src regulates TRPM8 function in an indirect manner. LCK has emerged as one of the key molecules to function in lymphocytes and stimulates several ion channels, especially Kv1.3 potassium channel (Desai et al, 2000; Kuras et al, 2012; Lepple-Wienhues et al, 1998; Lepple-Wienhues et al, 2001; Szigligeti et al, 2006). LCK coupled with hDlg indirectly regulated Kv1.3 channel activities (Hanada et al, 1997). Apart from Kv1.3, it is not clear that LCK stimulates other channels in a direct or indirect manner. Our data revealed that LCK directly interacted with TRPM8 by protein-protein interaction assay *in vitro* and positively modulated TRPM8 phosphotyrosine and ITRPM8 densities, which expands the mechanism of LCK function on ion channels. Notably, BLK or LYN interacted with TRPM8, but neither affected TRPM8 phosphotyrosine nor ITRPM8 densities. Thus it is interesting to assess the physiological role of TRPM8 interaction with BLK or LYN.

TRPM7, belonging to TRPM family with TRPM8, binds to 14-3-3, which requires autophosphorylation of TRPM7 at S1403 (Cai et al, 2018). However, 14-3-3 involved in regulating TRPM7 cellular localization (Cai et al, 2018), exhibiting clear difference to regulating TRPM8 multimerization. Additionally, our data revealed the importance of 14-3-3ζ for LCK and impaired phosphotyrosined TRPM8 (TRPM8-Y1022F) regulating the TRPM8 multimerization, supporting that TRPM8 phosphotyrosine modulated 14-3-3ζ mediated channel multimerization. Previous studies showed binding of 14-3-3 to proteins usually occurs after phosphorylation of Ser/Thr within two conserved consensus motifs (RSXpS/TXP or RXXXpS/TXP), giving rise to a variety of functional consequences for regulating its activation or deactivation (Gardino et al, 2006; Liu et al, 2021). Apart from phosphotyrosine, LCK as a tyrosine kinase, was also found to increase the level of TRPM8 phosphoserine. TRPM8-Y1022F markedly inhibited the level of TRPM8 phosphoserine, further supporting that TRPM8 phosphotyrosine affected its own phosphoserine level. Thus, we speculated TRPM8 phosphoserine might be involved in the process of 14-3-3ζ mediated channel multimerization regulated by LCK, although the involved exact phosphoserine site(s) remained to further study. Together, following phosphorylating TRPM8 at Y1022, LCK coordinately enhanced the TRPM8 phosphoserine for recruiting 14-3-3ζ and providing cross-bridging of 14-3-3ζ and TRPM8 in a complex, leading to TRPM8 multimerization for elevated I_TRPM8_ densities on the PM.

LCK as a Src family tyrosine kinase was originally identified to play an important role on T-cell functions (Adler et al, 1988; Voronova & Sefton, 1986). Now, LCK has been found to function as an oncogene in leukemia and various solid cancers, including breast cancer, colon cancer, and lung carcinoma (Bommhardt et al, 2019). Indeed, several LCK inhibitors have been to approve for the treatment of leukemia and various solid cancers including pancreatic cancer (Chen & Chen, 2015; Creeden et al, 2020; Lindauer & Hochhaus, 2014; Marech et al, 2014). Recently, Lck activity has been mainly regulated via phosphorylation/dephosphorylation of crucial tyrosine residues, Y394 and Y505 (Eck et al, 1994; Peri et al, 1993; Yamaguchi & Hendrickson, 1996). Our data revealed that TRPM8 phosphotyrosine feedback altered the level of phosphorylated LCK on Y505, but not Y394, which is responsible for the elevated activity of LCK. Therefore, TRPM8 phosphotyrosine suppressed the phosphorylation of LCK on Y505 in the inactive state thereby enhancing LCK activity. Several studies revealed that ubiquitination is also involved in the regulation of LCK activity in a diverse manner (Choi et al, 2010; Giannini & Bijlmakers, 2004; Rao et al, 2002). Heat shock protein 90 (Hsp90) prevents the active form Y505F of mutant LCK from being targeted for degradation by ubiquitination (Giannini & Bijlmakers, 2004). Apart from HSP90 mediated LCK ubiquitination without altering LCK expression, other regulators, such as Cbl and SOSC-6, modulated the degradation of LCK (Choi et al, 2010; Giannini & Bijlmakers, 2004; Rao et al, 2002). TRPM8 phosphorylation positively modulated LCK ubiquitination without affecting LCK expression, especially the ubiquitination of inactive form Y394F of mutant LCK. Thus, we elucidated the mechanism by which TRPM8 mediated ubiquitination at inactive LCK form regulating its kinase activity, which is obviously different from previous studies (Giannini & Bijlmakers, 2004). Nevertheless, how TRPM8 coordinates in the regulation of the crosstalk of phosphorylation and ubiquitination across LCK in different active states to modulate its activity needs to be further investigated.

In summary, we establish a link between LCK, 14-3-3ζ and TRPM8 and provide mechanistic insights into LCK-14-3-3ζ-TRPM8 axis for full understanding of TRPM8 multimerization mediated channel function and LCK activity. Through targeting inhibition LCK-14-3-3ζ-TRPM8 axis to impair oncogene function both of TRPM8 and LCK, it may enhance tumor sensitivity to therapeutics, which was utilized for a potential pharmacological use as a target for anti-cancer therapy.

## Materials and Methods

### Antibodies and reagents

A rabbit anti-LCK (#12477, PTGCN, China), anti-GFP (#50430, PTGCN), anti-14-3-3 (#14503,PTGCN), anti-Phosphotyrosine (p-Tyr) (Sigma-Aldrich), anti-Phosphothreonine (Cell Signaling Technology) Anti-Phosphoserine (Abcam, ab9332), anti-phospho-Lck-Y505 (pY505) (#MAB7500, R & D), anti-TRPM8 (#ACC-049, Alomone, Israel), anti-LC3B (#18725, PTGCN, China), anti-SQSTM1/p62 (#BM4385, Boster, China), anti-ULK1 (#20986, PTGCN), anti-phospho-ULK1 (Ser317) (#12753, Cell Signaling Technology), anti-AMPKa1 (#BM4202, Boster), and anti-phospho-AMPKa (Thr172) antibodies (#2535, Cell Signaling Technology), and mouse anti-phospho-Lck-Y394 (pY394) (#2751, Cell Signaling Technology) antibodies were used at a dilution factor of 1:1000. A mouse anti-GFP antibody (#66002, PTGCN), anti-HA (#M180, MBL, Japan), anti-Flag (#M185, MBL), anti-Myc (#M192, MBL), and anti-GAPDH (#60004, PTGCN) were used at a dilution factor of 1:3000. A goat anti-rabbit or anti-mouse HRP-conjugated secondary antibody obtained from Millipore were used at a dilution factor of 1:20000. The compounds for 4-amino-5-(4-chlorophenyl)-7-(dimethylethyl) pyrazolo[3,4-d] pyrimidine (PP2), sodium orthovanadate (Na3VO4), disuccinimidyl suberate (DSS), MG132, and saracatinib were obtained from Selleck. All reagents for cells culture were obtained from Invitrogen.

### Cells culture and transfection

Human cervical cancer cell line HeLa, human embryonic kidney 293T (HEK293T), and human pancreatic cancer cell lines PANC-1, AsPC-1 were obtained and cultured as previous described (Zhou et al, 2020). All cell lines were maintained in Dulbecco’s modified Essential medium (DMEM) containing 10% fetal bovine serum (FBS), L-glutamine (2 mM), penicillin G (100 units/ml) and streptomycin (10 mg/ml) at 37 °C with 5% CO_2_. Cells with 60~70% confluent were transfected with the indicated expression construct or siRNA using Lipofectamine™ 2000 Transfection Reagent (Invitrogen) according to the manufacturer’s instructions. After 48 hr of transfection, cells were harvested for follow-up corresponding experiments.

### Expression constructions and siRNA

The expression constructions for full-length rat Trpm8 (NM_134371) in pcDNA3 (pcDNA3-TRPM8), in pcDNA3.1-N-Flag (Flag-TRPM8), in pEGFP-N1 (GFP-TRPM8) were previously described (Zhou et al, 2020). The truncated expression constructs for GST fused C-terminus of TRPM8 (GST-M8C) in pGEX-4T-1, GFP fused C-terminus of TRPM8 (GFP-M8C) in pEGFP-C1, Flag-tagged cytosolic domains of TRPM8 (1-691 for M8-N, 756– 759 for M8-LI, 815–829 for M8-LII, and 980–1104 for M8-C) that subcloned into the pCMV10-3×Flag vector were previously described (Huang et al, 2020). The expression construct for mutant TRPM8 with mutation Y1022F were introduced using a PCR-based mutagenesis method (Huang et al, 2021; Huang et al, 2016; Huang et al, 2017). The human BLK, LCK, and LYN cDNAs were kindly provided by Pro. Jiahuai Han (Xiamen University, China) and subcloned into pcDNA3.1-HA to express HA fused BLK, LCK, and LYN in mammalian cells, respectively. LCK cDNA was also subcloned into pET28α(+) to express His fused LCK (His-LCK) in *E.coli* BL21. The human 14-3-3ζ cDNA from HEK293T cells was subcloned into pEGFP-C1 to express GFP fused 14-3-3ζ in mammalian cells. All expression constructs were verified by direct DNA sequencing analysis. The siRNA targeting human LCK (siLCK#1: 5’-UCAAGAACCUGAGCCGCAATT-3’ and siLCK#2: 5’-GGCAGCCCAAAUUGCAGAATT-3’), siRNA targeting human 14-3-3ζ (si14-3-3#1: 5’-GCCUGCAUGAAGUCUGUAATT-3’, si14-3-3#2: 5’-CGUCUCAAGUAUUGAACAATT-3’ and si14-3-3#3: 5’-CACGCUAAUAAUGCAAUUATT-3’), and scrambled control siRNA (5’-UUCUCCGAACGUGUCACGUTT-3’) were designed and synthesized by GenePharma (Suzhou, Jiangsu, China). siRNA knockdown efficiency was verified using western blotting analysis with anti-LCK antibody. The TRPM8 mRNA was detected by real time qRT-PCR using the following primers: Forward: 5’-TCTGCCGACCTTCAGGAGGT-3’, Reverse: 5’-ATGGAGTTCCACATCCAAGTCC-3’.

### Lentiviral production and creation of stable cell lines

The lentiviral production and creation of stable cell lines were performed as described previously (Huang et al, 2021; Huang et al, 2016; Huang et al, 2017). The DNA fragment of TRPM8 was subcloned into the lentiviral plasmid pCDH-CMV-MCS-EF1-turboRFP-T2A-Puro. HEK293T cells in 10-cm dishes were transfected with 5 μg of lentiviral constructs together with viral packaging plasmids 3 μg of psPAX2 and 3 μg of pMD2.G (All of related viral plasmids were kindly provided by Dr. Xiaorong Zhang (Institute of Biophysics, Chinese Academy of Sciences, China). After 48 h of transfection, the viral supernatant was harvested, filtered through a 0.22 μm filter, and then added to PANC-1 or AsPC-1 cells in 6-cm dishes with 10 μg/μl polybrene (Solarbio, H8761). At 48 h of viral infection, the cells were selected and cultured by replacing a new culture medium containing puromycin (Solarbio, IP1280) every 3-4 days for several weeks. The clones of puromycin-resistant cells stably expressing TRPM8 were isolated, characterized, and expanded in complete culture medium supplemented with 2 μg/ml puromycin.

### Western blot and immunoprecipitation

The western blot (WB) and immunoprecipitation were performed following the procedure described previously (Huang et al, 2016). Briefly, cells were lysed with lysis buffer (50 mM Tris-HCl, pH 7.5, 150 mM NaCl, 0.5% NP-40 and 1 mM EDTA supplemented with 1 × protease inhibitor complete mini EDTA-free cocktail from Roche). The supernatants of cell lysates were boiled for 5 min in 1 × SDS loading buffer (6 ×, 0.3 M Tris-HCl, 6% SDS, 60% glycerol, 120 mM dithiothreitol (DDT) and proprietary pink tracking dye), then subjected to 8-15% sodium dodecyl sulphate-polyacrylamide gel electrophoresis (SDS-PAGE) and transferred to polyvinylidene difluoride (PVDF) membrane. Blocking with 5% non-fat dry milk in TBST (20 mM Tris-HCl, 150 mM NaCl, 0.05% Tween-20), the membrane was then incubated with the indicated primary antibodies, secondary antibodies, and supersignal west pico plus (Invitrogen, 34,580) according to the manufacturer’s instructions. Finally, the protein signal on the membrane was recorded with a ChemiDoc XRS system (Bio-Rad Laboratories, Richmond, CA), and analyzed using the Image Lab software (Bio-Rad Laboratories).

For immunoprecipitation, 2 μg of the indicated antibody together with 500 μl of cell lysates (500 μg) were mixed at 4 °C for 3 h, followed by incubation with 30 μl of Protein-A/G beads (Santa Cruz Biotechnology) for 2 h. After three washes three times with lysis buffer supplemented with 0.1% Tween, the resulting immunocomplexes were subjected to WB assay. Studies were repeated at least three times.

For the ubiquitination assay, cells treatment with 10 μM concentration of the proteasome inhibitor, MG132, for 6 h were harvested and lysed with the denaturation buffer (6 M guanidine-HCl, 0.1 M Na_2_HPO_4_/NaH_2_PO_4_, 10 mM imidazole) as described previously (Huang et al, 2020). The supernatant of lysates was mixed with the indicated antibody, and then with Protein-A/G beads for 3 h with rotation at RT, followed by washes and WB assay.

### GST pull-down assay and protein-protein interaction Assay *in vitro*

The GST pull-down assay was perform as described previously (Huang et al, 2020). Recombinant proteins for GST-M8C or GST alone expressing in *E.coli* BL21 cells were purified using glutathione bead according to the manufacturer’s protocol (Thermo). Purified GST-M8C or GST alone were incubated with protein lysates extracted from HEK293T cells overexpressing HA-BLK, HA-LCK or HA-LYN, respectively. Four washes with the lysis buffer, the complex of proteins-bound GST-agarose bead was washed four times with sonication buffer (0.5% Nonidet P-40, 50 mM Tris/HCl, 150 mM NaCl, 1 mM EDTA supplemented with 1 × protease inhibitor cOmplete Mini EDTA-free mixture from Roche) and subjected to WB assay.

For protein-protein interaction assay *in vitro*, expression of His-LCK fusion protein in *E.coli* BL21 cells was induced with 1 mM isopropyl 1-thio-β-D-galactopyranoside (IPTG) for 8 h at 20 °C. The cells were then harvested, resuspended in sonication buffer and sonicated on ice. Following centrifugation, the supernatants were incubated with the Ni-NTA agarose (Beyotime, Shanghai, China) with rotation for 3h at 4 °C. The immobilized His-LCK was then washed with sonication buffer, 2 mM and 5 mM imidazole. Elution with 50 mM imidazole, 150 μg of bound His-LCK proteins were incubated with the above purified the complex of GST or GST-M8C proteins-bound GST-agarose bead for 1h at room temperature (RT). Washing with sonication buffer four times, the complex of proteins-bound GST-agarose bead were subjected to WB assay.

### Kinase assay *in vitro*

LCK kinase assay *in vitro* experiments was performed on a modified protocol as previously described (Moogk et al, 2016; Nika et al, 2010). LCK proteins extracted from HEK293T cells overexpressing HA-LCK were immunoprecipitated with anti-HA antibody. The immune-complex was extensively washed with lysis buffer twice, then washed with kinase assay buffer (20 mM Tris-HCl pH 7.5, 10 mM MgCl_2_, 10 mM MnCl_2_), and incubated together with 100 ng of bacterially purified GST-M8C in kinase assay buffer supplemented with 1 mM ATP. After incubation for 30 min at 37 °C for 30 min, the reaction mixtures were terminated and boiled for 5 min in 1 × SDS sample buffer, followed by WB assay to detect the phosphotyrosine of C-termini of TRPM8.

### Electrophysiological experiments

For electrophysiological experiments, whole-cell cation currents mediated by TRPM8 (ITRPM8) was recorded by whole-cell patch-clamp technologies with an Axon MultiClamp 700B amplifier using the Digidata1550A digitizer (Axon Instruments, Sunnyvale, CA) as described previously (Moogk et al, 2016; Nika et al, 2010). Briefly, the indicated expression constructs including pEGFP-N1 and Flag-TRPM8 were transfected into HEK293T cells. After 48 hr of transfection, we selected the cells expressing an approximately equal amount of GFPs for recording I_TRPM8_ at RT in an extracellular solution containing (mM) 150 NaCl, 6 CsCl, 1 MgCl_2_, 1.5 CaCl_2_, 10 glucose, 10 mM HEPES, pH 7.4 with NaOH. The peptides were filled with pipette solution ((mM): 150 NaCl, 3 MgCl_2_, 5 EGTA, 10 HEPES, pH 7.2 with NaOH) to form a tip resistance of 2 ~ 4 MΩ. Series resistance was compensated by 75–85% to reduce voltage errors. The holding potential was −60 mV and details of each pulse protocol are given schematically in the related figures. Densities of whole-cell TRPM8 current (ITRPM8) was normalized to the cell capacitance (pA/pF). The data were analyzed using a combination of Clampfit version 11.0 (Molecular Devices), Microsoft Excel, and GraphPad Prism 5 (GraphPad Software Inc., San Diego, CA, USA).

### Cell surface biotinylation assay

Isolation of PM proteins by cell surface biotinylation assay was described previously (Huang et al, 2021; Huang et al, 2016; Huang et al, 2017). Briefly, cells were harvested and incubated with sulfo-NHS-SS-Biotin to label the PM proteins in ice-cold PBS for 30 min, followed by incubating with 100 mM glycine to quench the biotinylation reaction. After three washes with PBS, cells were harvested in the above lysis buffer. The biotinylated proteins were precipitated with NeutrAvidin-agarose resins beads (Pierce) overnight at 4 °C. The proteins-beads complex were washed with lysis buffer, and then resuspended in 1 × SDS loading buffer for WB assay.

### Immunocytochemistry and confocal microscopy

Immunocytochemistry was performed as described previously (Huang et al, 2020; Zhou et al, 2020). Transfected HEK293T, HeLa or ASPC-1 cells on the glass coverslips were washed three times with ice-cold PBS, fixed for 10 min with 4% paraformaldehyde (w/v) in PBS, and permeabilized for 15 min by incubation with 0.2% Triton X-100 at RT. After blocking with blocked with 1× PBS containing 0.1% Triton X-100 (v/v) and 10% goat serum (v/v) for 2 h, the samples were incubated with the indicated primary antibodies (e.g., anti-Ki67 antibody (#27309, PTGCN)) overnight at 4°C and fluorescence-labeled secondary antibodies in PBS supplemented with 2% FBS and 1% BSA for 2 h at RT. DAPI (1 μg/ml, Solarbio, C0065) was used for nuclei staining. Finally, the samples were washed three times with ice-cold PBS and observed with a confocal laser-scanning microscopy (Leica SP8, Wetzlar, Germany). At least three fields of view were analyzed. Data analysis was performed using the Leica LAS AF Lite software.

### 5-Ethynyl-20-deoxyuridine (EdU) incorporation assay

The EdU incorporation assay was performed as described previously (Zhou et al, 2020). EdU labeled Transfected PANC-1 cells were examined using the BeyoClick™ EdU Cell Proliferation Kit with Alexa Fluor 488 (Beyotime, C0071S), and then imaged under an Olympus FSX100 microscope.

### *In Vitro* colony formation assa*y*

Colony formation assay was performed as described previously (Huang et al, 2020; Zhou et al, 2020). An approximately equal amount of AsPC-1 cells transfected with vector, wild type or mutant TRPM8 were seeded in 12-well plates and allowed to growth for 7 ~ 10 days. The medium was replaced every 3 days. Cells were washed twice with PBS, fixed with 4% paraformaldehyde, and stained with 0.5% crystal violet staining solution (Sigma-Aldrich, USA). Colonies with more than 50 cells in triplicate wells were counted.

### *In vitro* cell migration assay

The effects of TRPM8 on migration ability of cells were evaluated using wound-healing assay and transwell assay. For wound-healing assay, stably maintained PANC-1 cells with 80% confluent in 12-well plates were cultured for 24 h after the formation of a monolayer. The monolayer was scratched with the tip of a 10 μL pipette, washed by PBS to remove the cell fragments, followed by adding the conditioned medium. The wound healed for 12 h, and was imaged at the same wound location using an Olympus FSX100 microscope.

For transwell assay, about 5×10^4^ of transfected cells were digested and placed into the upper chamber precoated 8 μm pore transwell insert (Fisher Scientific, 0877121) with the lower chamber containing the medium (containing 10% FBS). After incubation for 24 hours at 37°C, 5% CO_2_. The upper surface of the membrane was then gently scraped using a cotton swab to remove the non-migrated cells and washed twice with PBS. The wells were then fixed in 4% paraformaldehyde for 30 min, permeabilized with 0.2% triton for 10 min, and stained with 0.5% crystal violet staining solution. Following washes twice with PBS, migrated cells were observed and photographed under an Olympus FSX100 microscope. The number of migrated cells was determined by averaging five random fields per well.

### Animal xenotransplantation engraftment experiments

Animal experiments in our study have been reviewed and approved by the use of laboratory animals by the Hubei University of Technology Animal Care and Use Committee. The zebrafish were maintained according to standard protocols (http://ZFIN.org) and embryos were grown at 28.5°C in egg water (60 μg/ml Instant Ocean sea salts). For zebrafish engraftment xenotransplantation, 2 days post-fertilization (dpf) embryos were utilized and obtained from adult AB zebrafish (Danio rerio). About 300 of RFP labeled stably maintained PANC-1 cells were inoculated in the blood circulation of 2 dpf zebrafish embryos as previously described (Tulotta et al, 2016). Prior to microinjection, the survival rate of cells was above 90% by capturing some cells for trypan blue staining and counting. Embryos were incubated at 34 °C for 4 days and imaged under anesthesia in egg water containing 200 μg/ml tricaine (Sigma Aldrich) using a fluorescence microscopy (Leica M205FA, Germany).

For xenografts engraftment in mice, BALB/c nude mice at 4–6 weeks of age (18-22 g) were utilized and purchased from Vital River Laboratory Animal Technology (Beijing, China). Three million stably maintained PANC-1 cells in 100 μl of phosphate-buffered saline (PBS) were subcutaneously implanted into the left and right axillae of female BALB/c nude mice per group by growth for 4-6 weeks as previously described (Zhou et al, 2020; Zhou et al, 2020). The tumor volume (V) was monitored and measured every 5 days with the following formula: V = [(tumor length × width × 2)/2], and the weight was calculated until the mice were sacrificed.

### Statistical analysis

Data are presented as mean ± SEM, and all data reported are based on at least three independent experiments. Student’s unpaired or paired two-tailed t tests (GraphPad) were carried out to determine statistical significance as appropriate. For comparisons of more than two groups, one-way analysis of variance was employed for normal distributions and the Kruskal-Wallis test for non-normal or small samples. *P* values less than 0.05 were considered statistically significant. *represents P < 0.05, **represents P < 0.01 and ***represents P < 0.001, NS stands for “not statistically different”.

## Acknowledgements

This study was supported by the China National Science Foundation (32070726 to JFT, 32000797 to YH), National Natural Science Foundation of Hubei (2020CFA073 to JFT), and the Wuhan Science and Technology Project (2019020701011475 to JFT). We also thank Pro. Qing Kenneth Wang for plasmid for wild-type Myc-Ub and Pro. Jiahuai Han for the human BLK, LCK, and LYN cDNAs.

## Author contributions

Y. Huang and S. Li write the main manuscript and performed molecular biology experiments. J. Tang, C. Zhou and Y. Huang designed the project and supervised all experiments. Z. Wang, S. Li, L Liu, W. Zhao, Q. Liu, K. Wang, D. Ali, M. Michalak, X.-Z. Chen provided support with experimental techniques and analyzed the data. All authors reviewed and approved the final manuscript.

## Expanded View Figure legends

**Figure EV1.**
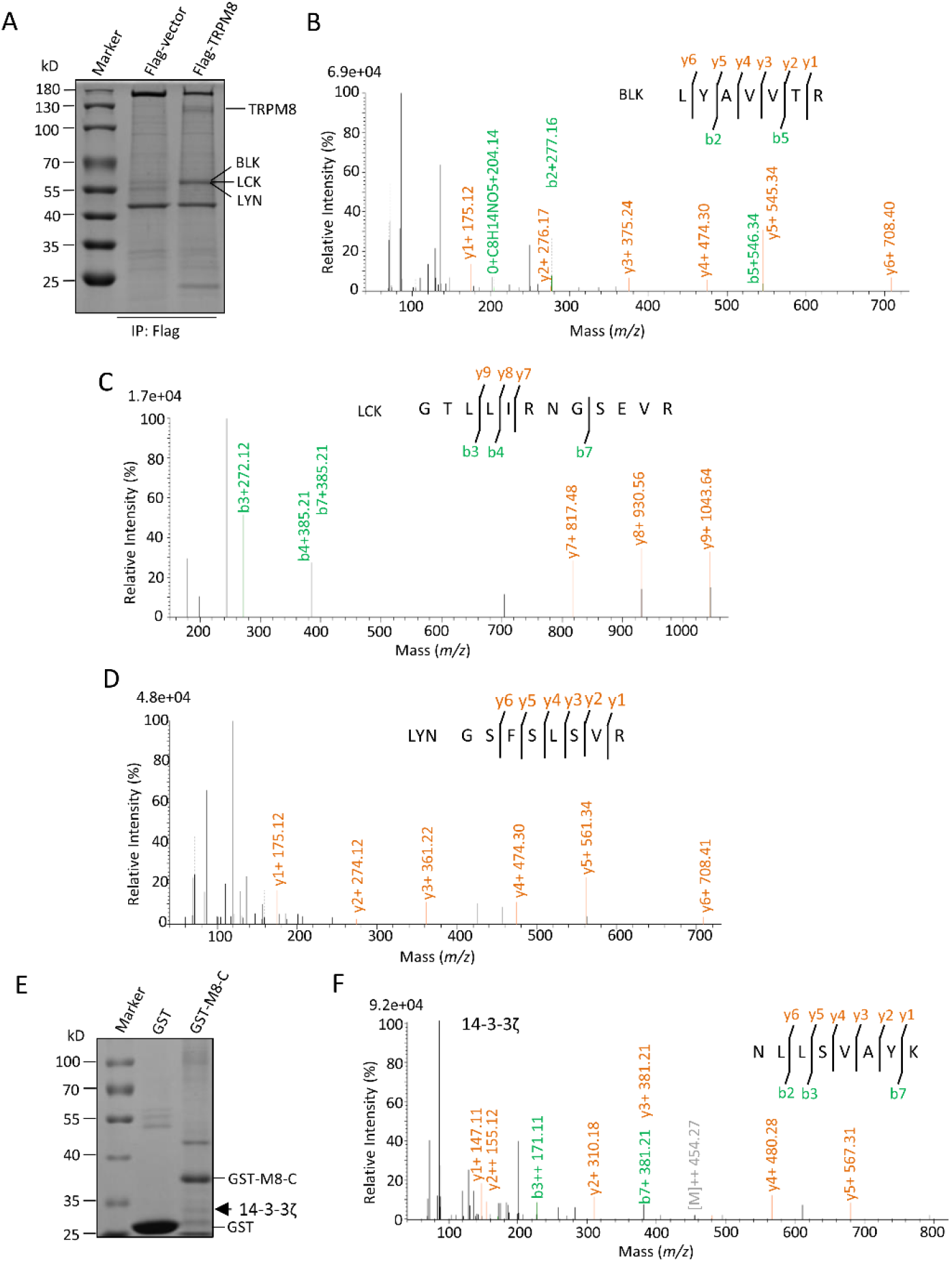
Identification of novel TRPM8-interacting partners by Co-IP assay. (**A**) Co-IP assay coupled with Coomassie staining. The lysates from MCF7 cells transfected with control vector or Flag-TRPM8 construct were precipitated with an anti-Flag antibody and subjected to Coomassie staining. (**B-D**) The relevant ~ 60 kD band (shown with an arrow) as in ***A*** was selected for mass spectrometric (MS) assay in combination with the NCBI Blast. Peptide sequences of MS assay are indicated in ***B*** (BLK), ***C*** (LCK), and ***D*** (LYN). (**E**) GST pull-down coupled with Coomassie staining. Purified GST alone or GST-M8C proteins expressing in *E.coli* BL21 bacteria were incubated with the lysates of MCF7 cells and subjected to Coomassie staining. The relevant band of ~ 30 kD protein (shown with an arrow) was selected for MS analysis. (**F**) MS imaging of 14−3−3ζ as a novel partner of TRPM8 in combination with the NCBI blast (peptide sequences are indicated).

**Figure EV2.**
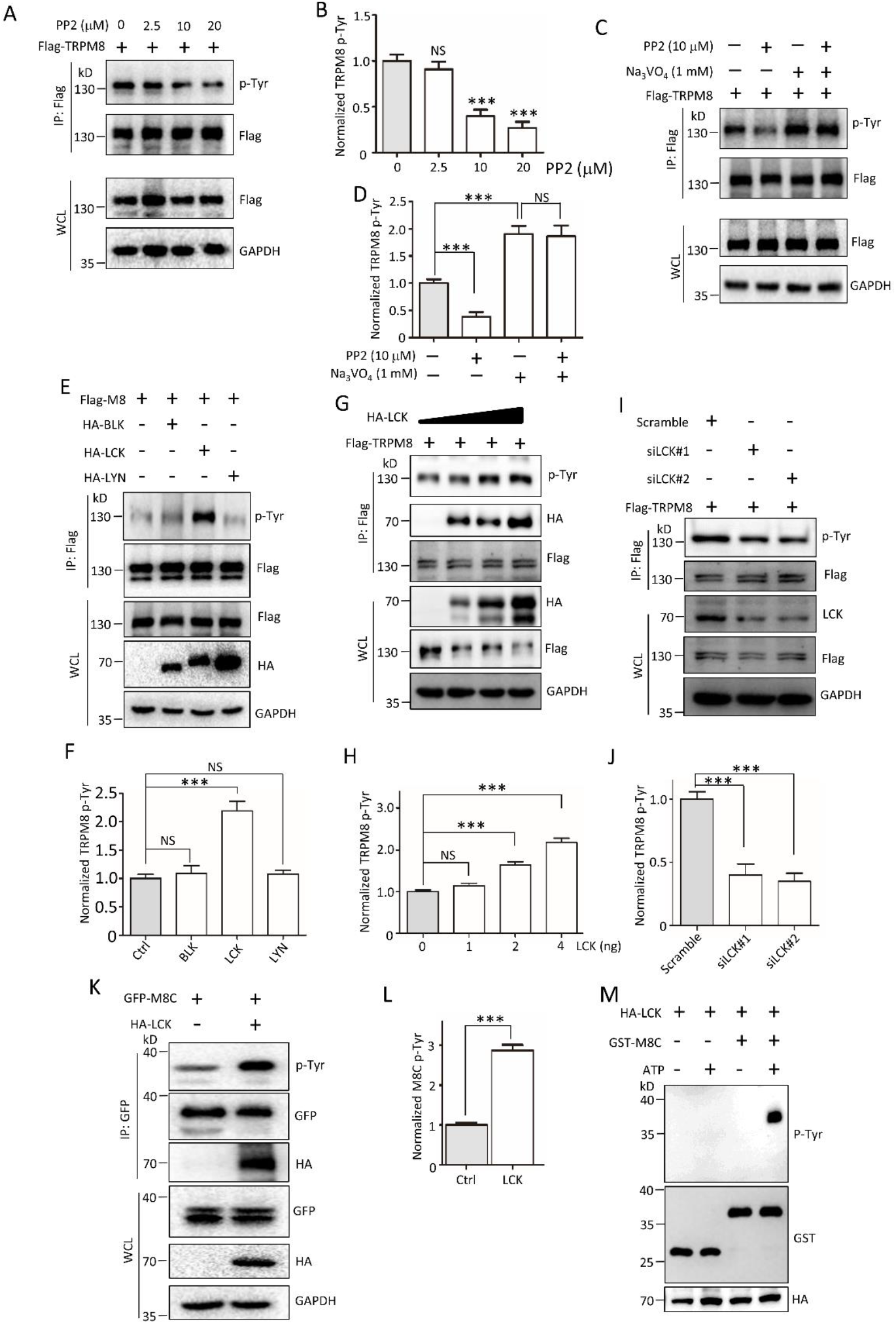
Effect of the compounds for PP2 and sodium orthovanadate, and LCK on TRPM8 phosphotyrosine. (**A-B**) HeLa cells expressing Flag-TRPM8 were incubated with DMSO dissolving different concentrations of PP2 (in μM, 0, 2.5, 10, 20) for 24 h before harvest, lysed and immunoprecipitated with an anti-Flag antibody. The samples were then analyzed by immunoblotting with the anti-Flag and p-Tyr antibodies to detect the level of TRPM8 phosphotyrosine. (**C-D**) Similar experiments in ***A*** and ***B*** but treatment with 10 μM PP2, 1 mM Na3VO4, or their combination. Na3VO4, sodium orthovanadate. PP2, 4-amino-5-(4-chlorophenyl)-7-(dimethylethyl) pyrazolo[3,4-d] pyrimidine, a Src family kinases inhibitor. (**E-F**) Expression constructs for Flag-TRPM8 were co-transfected into PANC-1 cells with HA-BLK, HA-LCK, or HA-LYN, respectively. The cells were then harvested for IP with an anti-Flag antibody and WB assay with the anti-Flag and p-Tyr antibodies to detect the level of TRPM8 phosphotyrosine. (**G-H**) Similar experiments in ***E*** and ***F*** but cells expressing with various amounts of HA-LCK. (**I-J**) Similar experiments in ***E*** and ***F*** but cells expressing with human LCK-specific siRNAs (siLCK#1 or #2) or negative scramble siRNAs. (**K-L**) HEK293T cells were co-transfected with GFP-M8C with HA-LCK or control vector, then harvested for IP with an anti-GFP antibodies, and WB assay with the anti-GFP and p-Tyr antibodies to detect the level of M8C phosphotyrosine. (**M**) Kinase assay *in vitro.* Purified GST alone or GST-M8C fusion proteins expressing in *E.coli* bacteria and HA-LCK immunoprecipitated with anti-HA antibody from HEK293T cells expressing HA-LCK constructs were mixed with or without 1 mM ATP in kinase assay buffer (20 mM Tris-HCl pH 7.5, 10 mM MgCl_2_, 10 mM MnCl_2_), and the reaction mixtures were then terminated, followed by WB assay with the indicated antibodies. ***, P < 0.001, NS, not significant. Data are presented as mean ± SEM. All studies were repeated at least three times.

**Figure EV3.**
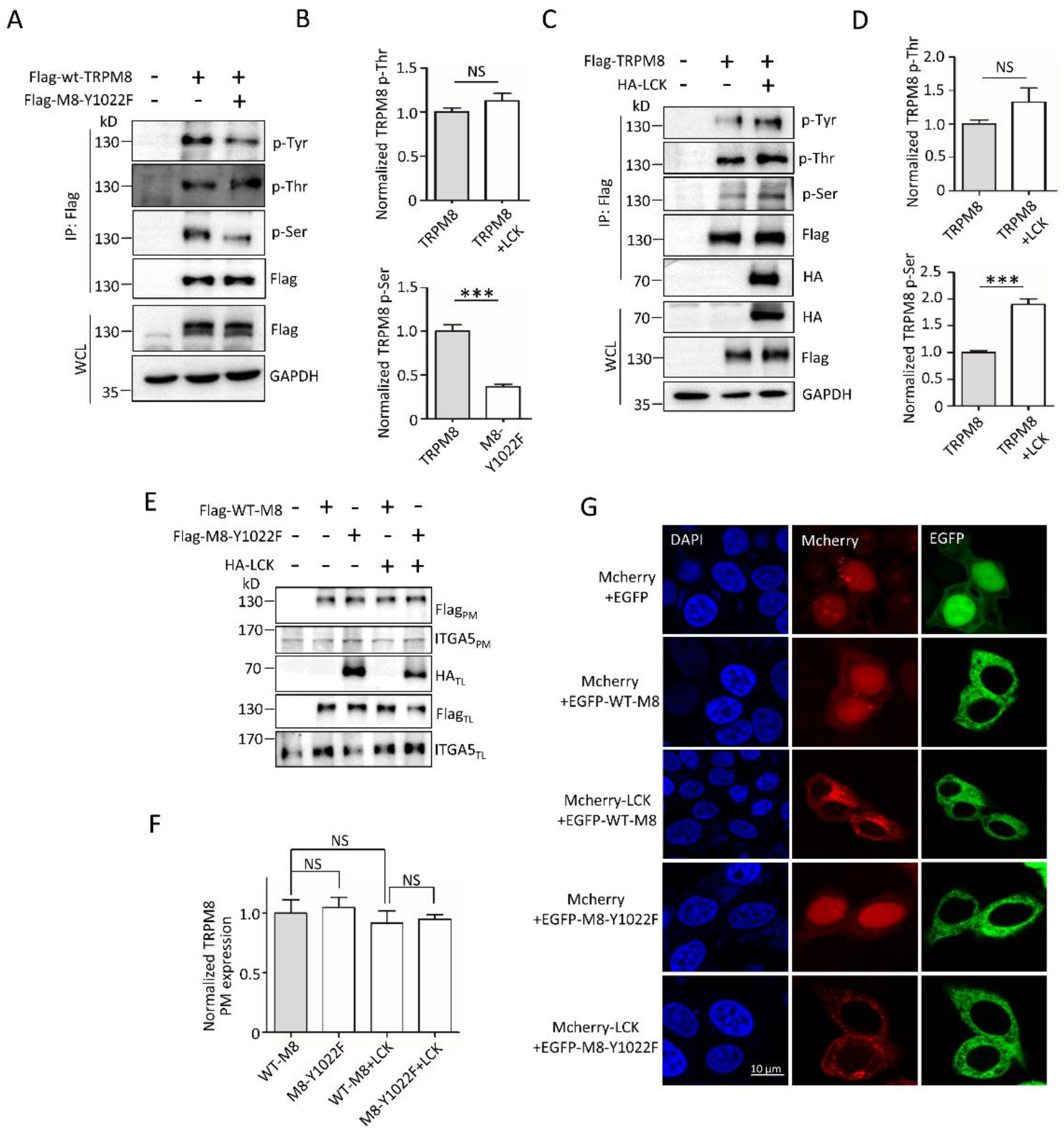
The effect of LCK or TRPM8-Y1022F on Ser/Thr phosphorylation of TRPM8 and LCK on mutant TRPM8-Y1022F expression on the PM. (**A-B**) HEK293T cells were transfected with Flag-WT-TRPM8 or Flag-M8-Y1022F, and harvested for IP with an anti-Flag antibody and WB assay with the indicated antibodies. (**C-D**) Flag-TRPM8 were co-transfected with or without HA-LCK into HEK293 cells. The cells were then harvested for IP with an anti-Flag antibody and WB assay with the indicated antibodies. (**E**) WB imaging of TRPM8 in the PM and total lysates from PANC-1 cells co-transfected Flag-WT-TRPM8 or Flag-TRPM8-Y1022F, in the presence or absence of HA-LCK. (**F**) Quantification of PM and total protein expression levels of TRPM8 in (***E***). (**G**) Representative confocal imaging of PANC-1 cells co-expressing mcherry-LCK with EGFP-WT-TRPM8 or EGFP-TRPM8-Y1022F. DAPI (1 μg/ml) was used for nuclei staining. ***, P < 0.001, NS, not significant. Data are presented as mean ± SEM. All studies were repeated at least three times.

**Figure EV4.**
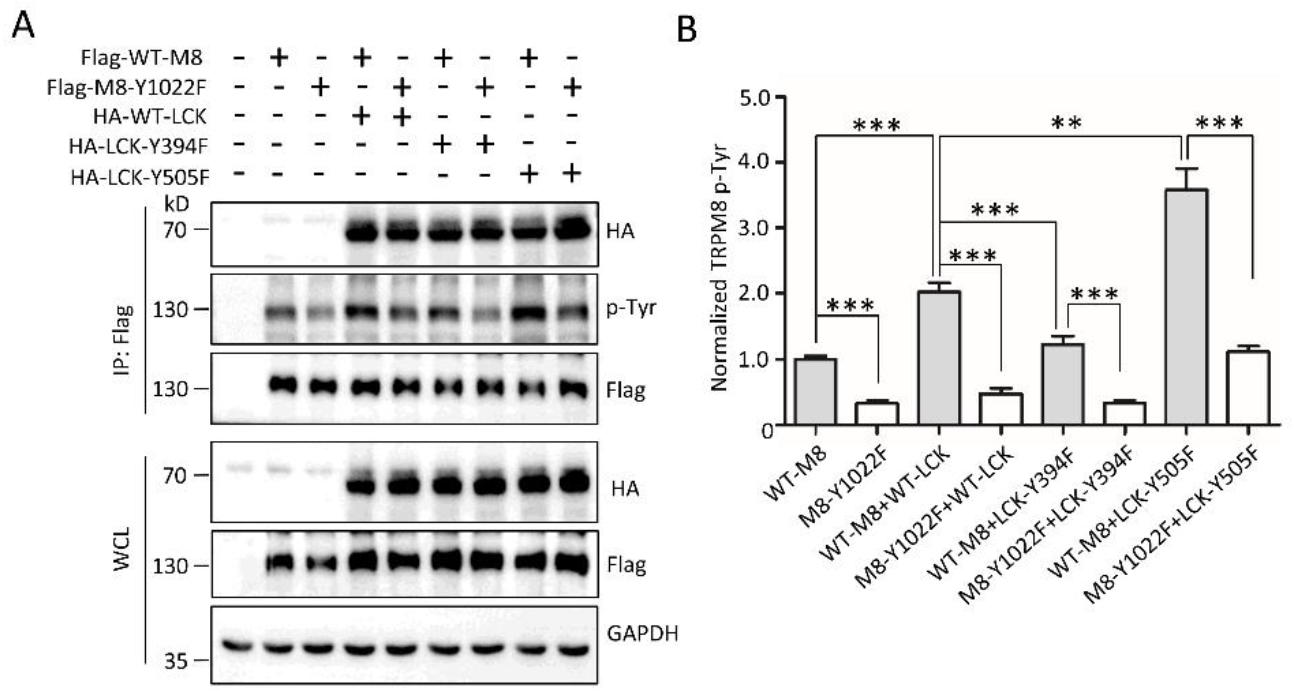
The effect of LCK mutants on TRPM8 phosphotyrosine. (**A-B**) Expression constructs for Flag-WT-M8 or Flag-M8-Y1022F were co-transfected with control vector, HA-tagged wild type, or mutant LCK into HEK293T cells. The cells were harvested for IP with an anti-Flag antibody and WB with the indicated antibodies to detect the level of TRPM8 phosphotyrosine. **, P < 0.01, ***, P < 0.001, NS, not significant. Data are presented as mean ± SEM. All studies were repeated at least three times.

**Figure EV5.**
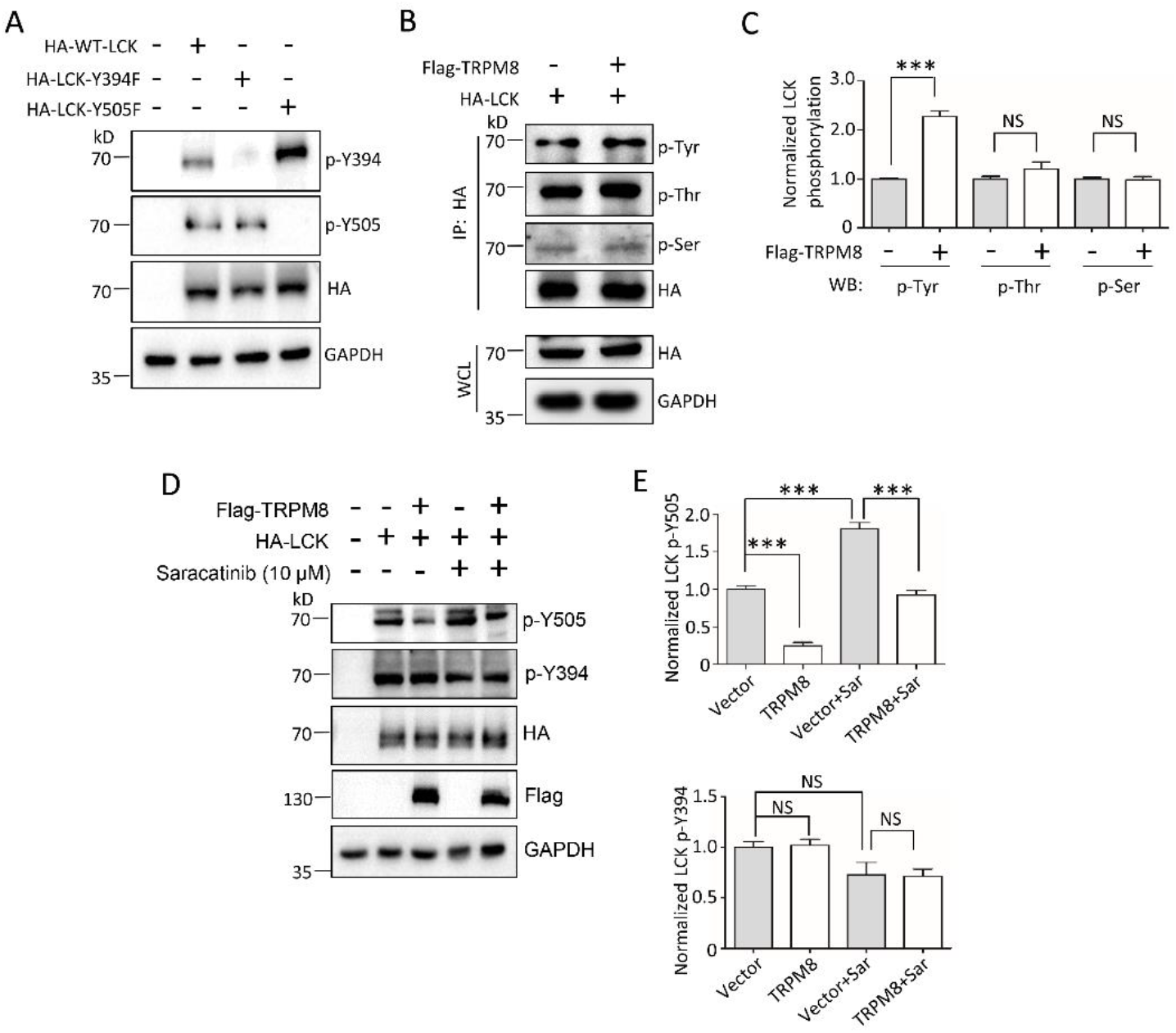
The effect of TRPM8 on LCK phosphorylation. (**A**) WB imaging of HEK293T cells individually transfected with control vector, HA-tagged wild type, or mutant LCK with the indicated antibodies. (**B-C**) PANC-1 cells were co-transfected HA-LCK with or without Flag-TRPM8, and then harvested for IP with an anti-HA antibody and WB assay with the indicated antibodies. (**D-E**) PANC-1 cells were co-transfected HA-LCK with or without Flag-TRPM8, before harvest for treatment with 10 μM saracatinib for 24 h. The lysates were subjected to WB assay with the indicated antibodies. ***, P < 0.001, NS, not significant. Data are presented as mean ± SEM. All studies were repeated at least three times.

